# Design and Synthesis of Triazine-Based Hydrogel for Combined Targeted Doxorubicin Delivery and PI3K Inhibition

**DOI:** 10.1101/2024.11.19.624181

**Authors:** Subhasis Mandal, Avinandan Bhoumick, Arpana Singh, Sukanya Konar, Arkajyoti Banerjee, Arnab Ghosh, Prosenjit Sen

**Author notes:** Corresponding Author: Prof. Prosenjit Sen, School of Biological Sciences, Indian Association for the Cultivation of Science (IACS), Kolkata-700032 West Bengal, India. Represent Equal contribution (Subhasis Mandal, Avinandan Bhoumick, Arpana Singh, Sukanya Konar, Arkajyoti Banerjee).

## Abstract

Melanoma, an aggressive skin cancer originating from melanocytes, presents substantial challenges due to its high metastatic potential and resistance to conventional therapies. Hydrogels, three-dimensional networks of hydrophilic polymers with high water-retention capacities, offer significant promise for controlled drug delivery applications. In this study, we report the synthesis and characterization of hydrogelators based on the triazine molecular scaffold, which self-assemble into fibrous networks conducive to hydrogel formation. Rheological analysis confirmed their hydrogelation properties, while microscopic techniques including FE-SEM and FEG-TEM provided insights into their morphological networks. The drug delivery capability of these hydrogelators was evaluated using doxorubicin, a widely employed anticancer agent, demonstrating enhanced biocompatibility and reduced side effects compared to free doxorubicin. Additionally, the hydrogelators exhibited inhibitory activity against phosphoinositide 3-kinase (PI3K), a key enzyme frequently mutated in cancer, and also involved in melanoma progression. The dual functionality of this delivery system – controlled drug release and PI3K inhibition – highlights the potential of triazine-based hydrogelators as innovative therapeutic platforms for melanoma treatment.

## INTRODUCTION

Melanoma, a malignant skin cancer originating from melanocytes, the pigment-producing cells, is strongly associated with excessive exposure to ultraviolet (UV) radiation from sunlight or artificial sources such as tanning beds. Over recent decades, the global incidence of melanoma has risen significantly, representing a growing public health challenge^1^. Early detection and timely treatment are essential to improving patient outcomes. While intravenous chemotherapy remains a standard treatment modality for many cancers, its effectiveness in melanoma is limited by systemic toxicity and poor tumor-specific targeting^2^. Researchers have explored local chemotherapy administration for certain cancers, including melanoma, which involves direct delivery of the drug to the targeted tumor site or a specific region of the body. This approach enhances drug efficacy by concentrating the therapeutic dose at the target site, minimizing systemic toxicity, and reducing off-target effects, thereby improving patient quality of life. Hydrogel-based systems have been used in clinical settings for effective drug delivery as they offer temporal and spatial control over the release of therapeutic agents, including cells, macromolecules, and small molecules^3^. They are highly adaptable, with tunable physical properties, controlled degradability, and the capacity to protect labile drugs from degradation. In cancer therapy, hydrogels have shown particular promise as carriers for chemotherapeutic agents, encapsulating drugs within their matrices to enable precise and efficient delivery^4^.

**Scheme 1.**
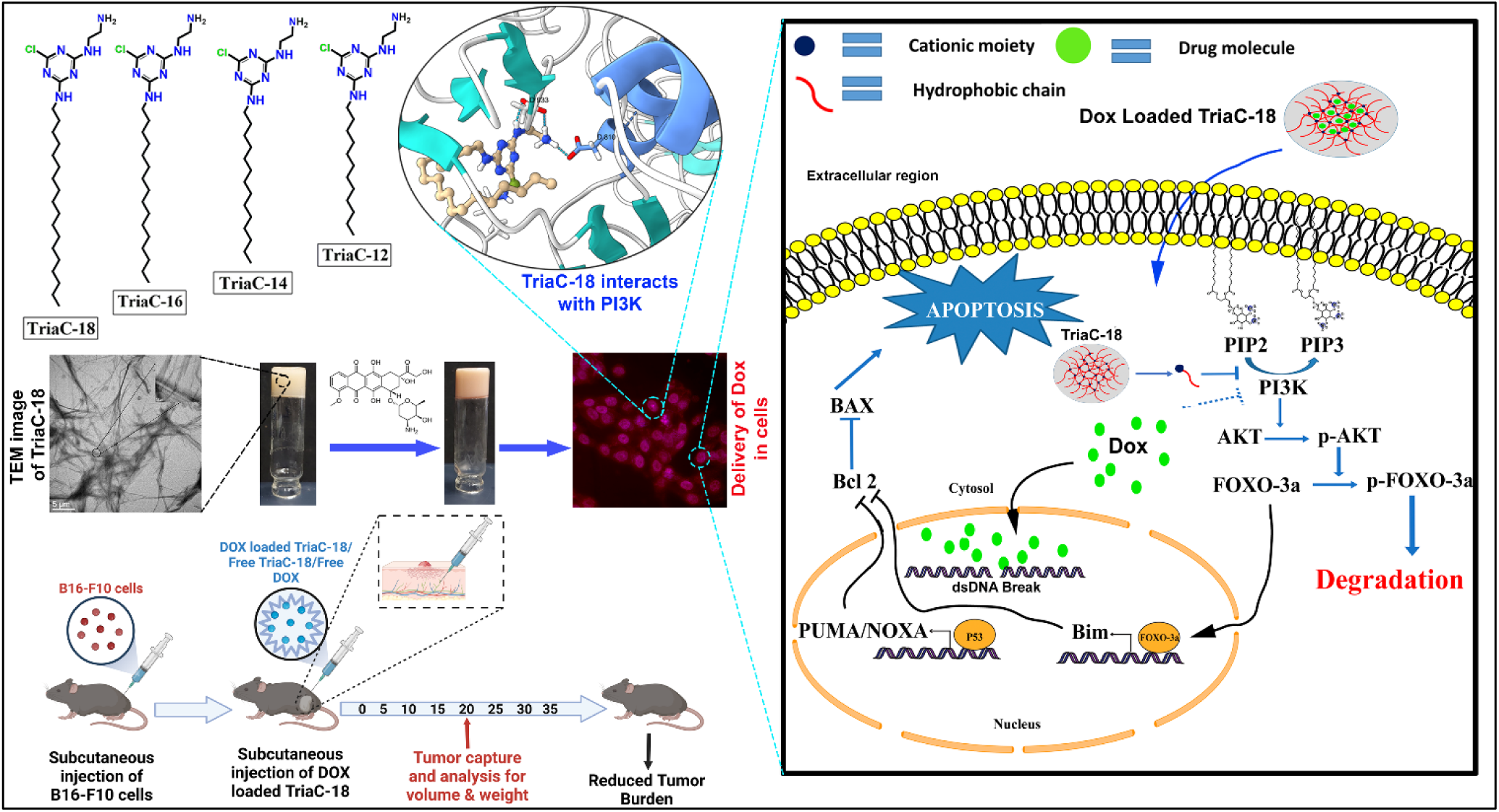
Exploring the Anticancer Potential of Triazine-Based Hydrogelators: Mechanistic Insights.

Hydrogels are three-dimensional networks of hydrophilic polymers capable of retaining 70–90% water, making them highly biocompatible and suitable for drug delivery applications^3^. While both natural and synthetic hydrogels have been widely studied, natural hydrogels are often preferred due to the potential toxicity associated with synthetic variants. Common examples of natural hydrogels include alginate^5^, gelatin^6^, fibrin^7^, polyethylene glycol (PEG)^8^, and hyaluronic acid-based hydrogels^9^. These hydrogels can respond to pathological pH changes, where ionization of acidic and basic groups in their networks induces swelling, enabling controlled drug release. Recent advances in hydrogel research have also highlighted their application in self-healing flexible sensors, emphasizing the importance of hydrogels with high stability, nontoxicity, biocompatibility, and robust mechanical properties for biomedical use^10^.

Triazine, a nitrogen-containing aromatic heterocyclic scaffold, is a ubiquitous structural motif in numerous drugs, such as antifungals, antibacterials, and cancer chemotherapeutics^11^. The presence of polar –N-H bonds enables them to interact with each other forming a supramolecular assembly. Different types of triazine-based hydrogels depending on their molecular scaffold have been reported so far like RADA16^12^ EAK16^13^ Fmoc-peptides^14^ ^15^ ^16^ peptide-amphiphiles^17^ and peptide-based polymers^18^. They can encapsulate and deliver a wide range of therapeutic agents, including proteins^19^ ^20^, nucleic acids^21^ ^22^, and small molecules^23^ ^22^. In the last two decades, triazine scaffolds have been employed to design inhibitors targeting proteins such as PI3K/m-TOR^24^ ^25^, anti-HIV activity^26^ ^27^, tubulin polymerization inhibitors, hCA IX inhibitors^28^, catalytic Topo-IIa inhibitor^29^, and others.

In this study, we synthesized a series of triazine-based hydrogelators with hydrophobic carbon chains of varying lengths: TriaC-12, TriaC-14, TriaC-16, and TriaC-18. Their gelation properties were confirmed through rheological analysis and morphological characterization using FE-SEM and FEG-TEM. Among these, TriaC-18 demonstrated superior hydrogelation, attributed to enhanced hydrophobic interactions and hydrogen bonding. Doxorubicin (DOX)^30^, a widely used chemotherapeutic agent, was effectively encapsulated within the TriaC-18 hydrogel. Cellular assays, including MTT, cell cycle analysis, nuclear condensation, Annexin V-FITC/PI staining, caspase-3/7 activity, and nuclear localization studies, revealed significantly enhanced anticancer activity of DOX-loaded TriaC-18 compared to free DOX in A375 (human melanoma) and B16-F10 (murine melanoma) cells. Interestingly, TriaC-18 also exhibited intrinsic anticancer properties, functioning as a PI3K inhibitor, which acted synergistically with DOX to enhance therapeutic efficacy. Expression analysis of pro-apoptotic and anti-apoptotic proteins confirmed increased apoptosis activation with DOX-loaded TriaC-18 treatment. These findings position TriaC-18 not only as a promising drug delivery carrier but also as an anticancer agent itself, offering dual therapeutic potential for sustainable and targeted melanoma treatment.

## EXPERIMENTAL SECTION

### General

All the reagents and solvents used here were for analysis or synthesis grade. 500 MHz and 400 MHz spectrometers were used to record ^1^H and ^13^C NMR spectra by Bruker spectrometer. The molecular weights of the synthesized compounds were determined by the MALDI-TOF instrument. SEM was recorded by JEOL-JSM-7500F field-emission scanning electron microscopy. TEM images were captured by JEOL-TEM instruments. Chemical shifts are reported in ppm with the solvent residual peak as a reference; DMSO-d_6_ (δ_H_ 2.5, 3.34, δ_C_39.85). The reaction was monitored by thin-layer chromatography (TLC) on silica-plated (silica gel 60-120, Merck) aluminum sheets, and spots were detected by UV (254 and 365 nm). Flash chromatography was performed using Merck Silica Gel 60-120. Absorbance was measured using a SpectraMax spectrophotometer. Dried hexane, toluene, diethyl ether, THF, and DMF were used. All other chemicals were used without further purification.

### Materials

Phosphate-buffered saline (PBS) and 3-(4,5-dimethylthiazol-2-yl)-2,5-diphenyltetrazolium bromide (MTT) were purchased from Sisco Research Laboratories Pvt. Ltd. (SRL). Cyanuric chloride, octadecylamine, hexadecylamine, tetradecylamine, dodecylamine, ethylenediamine, di-tert-butyl decarbonate, and sodium bicarbonate were purchased from TCI (Tokyo Chemical Industry Co. Ltd.).

### Synthesis

All the characterization data of the molecules (^1^H NMR, ^13^C NMR, MALDI-TOF) have been provided in the Supporting Information.

#### Step I

A stirred solution of cyanuric chloride (1.93 g, 10.5 mmol) in acetone (500 mL) was cooled at 0°C, and a solution of appropriate amine (8.76 mmol) was added for ∼30 min dropwise in this solution. The mixture was stirred at 0°C for 2 h, TLC was checked, and after the completion of the reaction the solvent was evaporated to provide a crude product. The crude product was purified by column chromatography over silica gel using 5% ethyl acetate/hexane as eluent.

#### Step II

To product I, (1 g, 2.3 mmol), Boc-protected EDA (0.368 g, 2.3 mmol) was added. NaHCO_3_ solution was added dropwise to the reaction mixture and was stirred for 2 h at 60°C and TLC was checked. After the completion of the reaction the product was purified with 50% Hexane: Ethyl acetate.

#### Step III

Product II was treated with TFA for 2 h and excess TFA was removed by rotary evaporator.

#### TriaC-18

^1^H NMR (400 MHz, DMSO-d_6_) δ 6.75, 3.26, 3.24, 3.23, 3.17, 3.08, 3.07, 3.05, 3.04, 2.94, 1.45, 1.37, 1.22, 0.85, 0.84, 0.82.

^13^C NMR (DMSO-d_6_) δ 168.61, 168.54, 167.99, 166.17, 165.94, 165.73, 165.37, 156.04, 155.97, 78.03, 77.97, 40.70, 40.53, 40.48, 40.36, 40.22, 40.08, 39.94, 39.80, 39.67, 39.53, 31.78, 29.51, 29.33, 29.25, 29.20, 29.06, 28.66, 28.62, 26.85, 26.78, 26.72, 22.57, 14.38.

#### TriaC-16

^1^H NMR (400 MHz, DMSO-d_6_) δ 6.80, 3.24, 3.23, 3.17, 3.08, 3.07, 3.05, 2.93, 1.45, 1.36, 1.23, 0.86, 0.84, 0.83.

^13^C NMR (DMSO-d_6_) δ 167.99, 165.92, 165.72, 156.04, 155.98, 78.05, 78.01, 40.68, 40.51, 40.46, 40.34, 40.20, 40.06, 39.93, 39.79, 39.65, 39.51, 31.77, 29.50, 29.45, 29.32, 29.23, 29.19, 29.05, 28.67, 28.63, 26.83, 26.75, 22.57, 14.41.

#### TriaC-14

^1^H NMR (400 MHz, DMSO-d_6_) δ 6.81, 3.24, 3.23, 3.17, 3.08, 3.07, 3.05, 3.04, 2.93, 1.47, 1.46, 1.37, 1.25, 0.85, 0.83.

^13^C NMR (DMSO-d_6_) δ 168.00, 165.95, 165.75, 156.06, 78.07, 40.59, 40.38, 40.17, 39.96, 39.75, 39.55, 39.34, 31.67, 29.11, 28.68, 26.76, 22.54, 14.40.

#### TriaC-12

^1^H NMR (400 MHz, DMSO-d_6_) δ 6.79, 3.24, 3.23, 3.17, 3.08, 3.07, 3.05, 3.04, 2.93, 1.47, 1.45, 1.36, 1.24, 0.85, 0.84, 0.83.

^13^C NMR (DMSO-d_6_) δ 168.61, 168.55, 167.99, 166.18, 165.94, 165.74, 165.53, 165.37, 156.03, 78.08, 40.71, 40.53, 40.32, 40.11, 39.90, 39.69, 39.48, 39.27, 31.67, 29.31, 29.12, 29.10, 28.66, 26.82, 26.75, 26.69, 22.53, 14.37.

### Preparation of the Hydrogels

0.15 mM of each hydrogelator were weighed into four separate screw-capped glass vials, followed by the addition of 1 mL of ultrapure water (Milli-Q) to each vial. The vials were then heated on a hot plate until the hydrogelator molecules were completely dissolved. Subsequently, the vials were cooled in a water bath for 5 minutes. Gelation occurred instantly upon cooling, and complete gelation was confirmed using the glass inversion technique.

### Rheological Studies

To explore the viscoelastic behaviour and the mechanical strength of the hydrogelators, Rheology studies were performed by using an Anton Paar, MCR 302. The experimental setup involved using parallel plates with a diameter of 25 mm and a 1 mm gap between them.

### TEM Sample Preparation

The sample for TEM was prepared by smearing the hydrogel at MGC on a carbon-coated Cu (300 mesh) TEM grid. The grid was dried under vacuum at room temperature for 1 day and used for recording TEM images using an accelerating voltage of 100 kV without staining. Then TEM images were recorded in a JEOL instrument.

### SEM Sample Preparation

The SEM sample was prepared by smearing the hydrogel at MGC on an SEM stub. The grid was dried under vacuum at room temperature for 1 day and used for recording SEM images. SEM images were recorded in JEOL instrument JEOL (Model: JSM-7500F).

### DOX loading capacity and Release Study

The drug loading was done by submerging hydrogel TriaC-18 into a 1 mL solution that contains the DOX. The drug-loading process takes up to 96 hours. The residual drug solution is measured using UV-VIS. The drug loading (%) is calculated using the equation below:

Drug loading (%) =(Ci-Cr)/Ci ×100%, where Ci is the mol amount of the chemical initially in the solution and Cr is the remaining mol amount of the chemical. The absorbance was measured using a Spectromax M2e UV Visible Spectrophotometer. The wavelength of maximum absorbance of the DOX in water is 496 nm.

The release of DOX has been scanned in two different buffer solutions: PBS (pH = 7.4) and acetate buffer (pH = 5.5). Two ml of each PBS and acetate buffer was placed separately on the top of DOX-loaded gel samples and incubated at 37 °C for several hours. At each time point (2, 5, 10, 24, 48 h), 200 μl of supernatant was taken out from the vials, and equal amounts of respective buffer solutions were added. The release has been studied from UV−vis analysis. These release profiles have been investigated in triplicates.

#### Cell Culture and bioimaging

The human melanoma cancer cell lines, A375 and the murine melanoma cancer cell line B16-F10 were obtained from ATCC and cultured in standard DMEM (HiMedia, India). The media was supplemented with 10% FBS and 1% penicillin-streptomycin (Invitrogen). Cells were maintained at 37℃ and 5% CO_2_ level. Cells were seeded and allowed to grow to confluence. Cells were treated with 1.4µg/ml DOX, 10.7µg/ml TriaC-18, and the DOX-loaded TriaC-18 for stipulated periods.

#### Cell viability Assay

A375 and B16-F10 cells were seeded in 96-well plates at a density of 0.01×10^6^ cells per well, each well containing 0.2 ml of complete media. After overnight incubation, different concentrations of DOX, TriaC-18, and DOX-loaded TriaC-18 were treated for 48 hours. Subsequently, 3-(4,5-dimethylthiazole-2-yl)-2,5 diphenyltetrazolium bromide (MTT) at a concentration of 1 mg/ml was added to each sample and incubated for 4 hours. After incubation, the supernatant was carefully removed, and 100µl of dimethyl sulfoxide (DMSO) was added to each well to dissolve the formazan crystals. The absorbance values at 570 nm were measured using a Microplate reader. To ensure reproducibility, the experiment was repeated at least three times. Statistical analysis was performed using GraphPad Prism 8 to determine the significance of differences between the various groups.

#### Cell Migration assay

A375 cells were seeded and allowed to grow until reaching 80%–90% confluence as a monolayer. Subsequently, a gentle mechanical scratch was created using a micropipette tip. Following this, the cells were treated with three different substances: 1.4µg/ml DOX, 10.7µg/ml TriaC-18, and DOX-loaded TriaC-18. Images of the cells were captured at three specific time points: 0 hours (immediately after wounding), 24 hours, and 48 hours after the scratch was made, utilizing an inverted fluorescence microscope. This experimental setup aimed to investigate the effects of the various treatments on cell migration and wound healing dynamics.

#### Colony formation assay

The colony formation assay was performed to assess the prolonged impact of DOX-loaded TriaC-18 on the colony-forming ability of human melanoma cancer cells (A375). Initially, A375 cells were seeded and allowed to incubate for 14 days. After the incubation period, the human melanoma cancer cells (A375) were treated with 1.4µg/ml DOX, 10.7µg/ml TriaC-18, and DOX-loaded TriaC-18. Following the treatment, the cells were allowed to grow in culture media, with regular changes of fresh media every 2 days for a duration of up to 14 days. On the 14th day, the cells were fixed in ice-cold 70% methanol and subsequently stained with 0.1% crystal violet (diluted in methanol). The colonies obtained were counted using Fiji software, with the colonies of the vehicle control group considered 100%. The results presented include a representative image and data collected from three independent experiments. This colony formation assay provided valuable insights into the long-term effects of DOX-loaded TriaC-18 on the colony-forming ability of A375 cells, contributing to the understanding of its potential impact on cancer cell growth and survival.

#### Cell Cycle assay

The impact of 1.4µg/ml DOX, 10.7µg/ml TriaC-18, and DOX-loaded TriaC-18 samples on the cell cycle of A375 cells was assessed using a flow cytometer. Following 24 hours of treatment, the cells were harvested and fixed with 70% cold ethanol, after which they were incubated overnight at −20°C. Subsequently, the cells were washed and suspended in phosphate-buffered saline (PBS). To evaluate the DNA content, the cells were stained with 200µl of propidium iodide/ribonuclease A (RNase A) for 1 hour at 37 °C. Finally, the DNA content was analyzed using a flow cytometer (BD FACS Aria III) to determine any alterations in the cell cycle caused by the treatment with DOX, TriaC-18, or DOX-loaded TriaC-18 samples.

#### Nuclear condensation assay

Following a 24-hour incubation period, the culture medium containing either DOX, TriaC-18, or DOX-loaded TriaC-18 was aseptically aspirated and replaced with fresh DMEM supplemented with Hoechst 33342 at a final concentration of 10 μg/mL each. This modification allowed for robust nuclear staining. Subsequently, the cells were incubated for an additional 15 minutes at a temperature of 37 °C to facilitate optimal dye uptake. Following this incubation period, the culture plates were meticulously rinsed with fresh phosphate-buffered saline (PBS) to eliminate any surplus dye. Subsequently, 1 mL of fresh PBS was gently added to each well to maintain cell integrity. After these preparatory steps, high-resolution fluorescence images of the cellular samples were captured using a fluorescence microscope. This methodology enabled the precise assessment of nuclear condensation, a pivotal parameter under investigation in our study.

#### Annexin V-PI assay

A375 cells were seeded and allowed to reach confluence, afterward, the cells were treated with 1.4µg/ml DOX, 10.7µg/ml TriaC-18, and DOX-loaded TriaC-18 for 24 hours. Subsequently, the cells were collected, washed with PBS, and then dual-stained using an Annexin V-FITC/PI detection kit from Sigma, following the manufacturer’s protocols. The stained cells were promptly analyzed using a flow cytometer (BD-FACS Aria III) to assess the effects of the treatments.

#### Caspase 3/7 activity assay

The caspase 3/7 activity assay was carried out using the caspase 3/7 activity assay kit from Sigma, following the manufacturer’s protocol. Initially, A375 cells were treated with 1.4µg/ml DOX, 10.7µg/ml TriaC-18, and DOX-loaded TriaC-18 for 24 hours. After the incubation, the cells were lysed using a Lysis buffer, and the resulting supernatant was transferred to a 96-well plate. Subsequently, 50µl of reaction buffer (containing 10mM DTT, Abcam) was added to each well, and then Acetyl-Asp-Glu-Val-Asp p-nitroanilide (Ac-DEVD-p-NA) substrate (200µl) was introduced into the wells. The plate was incubated at 37°C and 5% CO_2_ for 90 minutes. After incubation, the absorbance reading was measured using a Microplate Reader at wavelength of 405nm. This assay enabled the quantification of caspase 3/7 activity in the treated A375 cells, providing valuable insights into the apoptotic response triggered by the different treatments.

#### Western blotting

The cells were lysed using 2X Laemmli buffer and heated at 95°C for 5 minutes. The proteins were separated by SDS-PAGE and transferred onto PVDF membranes. Subsequently, the membranes were blocked with 5% BSA in TBS for 1 hour and then incubated overnight at 4°C with primary antibodies specific to the proteins of interest (Supplementary Table 1). After washing the membranes with TBS-Tween 20 (0.1%), secondary HRP-tagged antibodies were added and incubated for one hour. Following this, the membranes were washed again, and the detection of protein bands was performed using the standard ECL method.

### Theoretical Calculations

#### Molecular Docking

##### Preparation of the protein and ligand

The crystal structure of the PI3Kα catalytic subunit p110α, bound to a triazine-based inhibitor PQR530 with the PDB ID “6OAC”^31^ was retrieved from RCSB Protein Data Bank^32^, and the protein structure was analyzed using PyMol. The co-crystallized ligand PQR530 (PDB ID: M1J) was bound to the PI3K catalytic subunit p110α active site in the kinase hinge region. The water molecules and co-crystallized ligand M1J were removed from the PDB file manually, and the missing residues were modeled using MODELLER^33^. The model was then validated with the help of the ERRAT score^34^ and the Ramachandran plot generated using the UCLA SAVES server. The modeled structure was then energy minimized using the Chiron Protein Structure Refinement server^35^. Subsequently, the prepared protein was then selected for the molecular docking of the ligand.

### Molecular Docking

The receptor and the ligand structure generated in ChemAxon Marvin JS were prepared in AutoDock Tools^36^by adding the partial charges. Docking was performed using AutoDock4.2^37^. A cubical receptor grid box of size 40 Å centered on the position of the co-crystallized ligand (M1J) with a search space of 0.5 Å was specified for the docking. Docking was then performed following Lamarckian GA with the default parameters. The docking results including the interactions were visualized and analyzed using PyMol, Discovery Studio Visualizer, and UCSF ChimeraX^38^. The 3D interaction profile was obtained using the PLIP (Protein-Ligand Interaction Profiler) server^39^.

### Molecular Dynamics Simulation

The starting configuration of the protein for the molecular dynamics simulation was selected based on the docking conformation. First, the ligand-docked configuration of the protein was kept at the center of a cubic simulation box of length 120 Å and then solvated with TIP3P water^40^. NaCl was added to neutralize the net charge of the system. The selected ligand (TriaC-18, abbrv. C18) was represented by the CHARMM CGenFF force field, obtained from the online server CGenFF^41^. the potential parameters of the protein and ions were obtained from the CHARMM36 force field with the potential form as below^42^ ^43^.

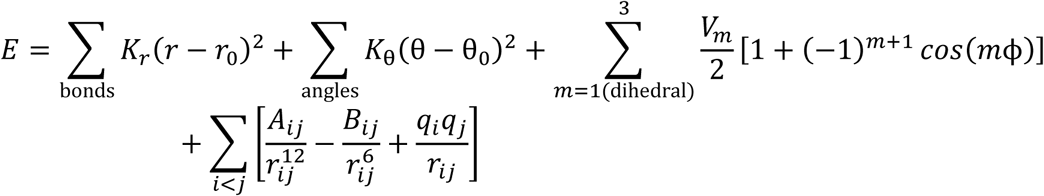

In detail, the first three terms represent the empirical interaction functions, namely, harmonic bonds, valence angles, and dihedral angles, respectively. Whereas, the last term represents the short-range 12-6 Lennard-Jones (LJ) potential between two interacting atoms i and j along with the long-range Coulomb potential. All atomistic simulations were performed using NAMD 2.12 simulation tools^44^. at first, we used the conjugate gradient method for 1000 steps to remove all energetically unfavorable contacts. After energy minimization, the simulation box was equilibrated for 125 ps in the NVT ensemble at 310 K temperature, applying harmonic restraint with a force constant of 10.0 for the backbone of the kinase and 5.0 for the side chain atoms excluding the hydrogen atoms. During the equilibration process, the ligand was fixed with a constraint scaling factor of 1.0. The temperature of the systems was kept constant at the desired temperature by using the damping coefficient (γ) of 1.0 ps-1 by Langevin dynamics. Long-range interactions were handled by the Particle Mesh Ewald (PME) method, with a real space cut-off of 12 Å and a 2 Å pair-list cut-off^45^ ^46^ ^47^. We used a 1-4 scaling factor in all our simulations. After that, the simulation box was equilibrated for another 250 ps in the NVT ensemble at 310 K temperature. In this equilibration process, the ligand was fixed with a constraint scaling factor of 1.0 and harmonic restraint with a force constant of 5.0 for the protein’s backbone and 2.0 for the side chain atoms excluding the hydrogen atoms. Then, the systems were equilibrated for 250 ps in the NPT ensemble to fix the simulation box length. In this last step of equilibration, the constraint scaling factor of the ligand was set to 0.5 and the harmonic restraint force constant of the protein’s backbone and the side chain atoms was set to 2.5 and 0.5, respectively. In the NPT simulations, the Noose-Hoover thermostat and barostat coupling constants were taken to be 0.05 ps and 0.025 ps respectively^48^ ^49^. The pressure was kept constant at 1.0 atm. The velocity Verlet algorithm with a time step of 1 fs was used^50^ for the production run. To analyze the different properties of the systems, for all cases, a production run of 250 ns was performed in the NPT ensemble at 310 K temperature. The trajectories were stored at an interval of 50.0 ps during the production run.

### Mice experiment

All animal experiments were conducted in compliance with the approved protocol by the Institutional Animal Ethics Committee. B16-F10 cells (5×10^5^) were implanted into the right or left flank of 6-week-old C57BL/6 mice. A single dose of 3.3 wt% TriaC-18 was administered subcutaneously at the tumor site for one group, 2.5mg/kg DOX injection subcutaneously at the tumor site for another group, and a third group received a subcutaneous injection of DOX-loaded TriaC-18 at the tumor site. The mice were continually monitored for 35 days, during which their body weights were recorded. After 20 days, the mice were euthanized, and both tumor weight and volume were measured. Tumor volume was calculated using the formula: V = (Length × Width^2^)/2.

### Statistical analysis

The experiments were conducted and repeated a minimum of three times. The data obtained were presented as the mean value ± standard deviation (SD) and analyzed using GraphPad Prism 8.0 software. The significance of the difference between different groups was determined by the Student’s t-test. A p-value cutoff of 0.05 was considered for statistical significance.

## RESULTS AND DISCUSSION

### Hydrogelation

Four hydrogelators were synthesized using triazine scaffolded as mention in Figure 1A. The self-assembly attributes, incorporating hydrophobic interaction^51^, π–π stacking^52^, and hydrogen bonding^53^, observed in these hydrogelators prompted exploration into the potential formation of opaque white hydrogels (TriaC-12, TriaC-14, TriaC-16, and TriaC-18) (Figure 1B). Frequency sweep experiments revealed their viscoelastic properties, characterized by the dominance of the elastic modulus (G’) over the loss modulus (G”) with minimal frequency (ω s⁻¹) dependence, particularly over extended durations (Figure 1C). The complex viscosity of the hydrogelators was also measured (Figure S17), and minimum gelation concentrations (MGC) were calculated (Figure 1D). Analysis of tan δ values provided insights into the viscoelastic behavior of the hydrogels (Figure 1E). Low tan δ values indicated robust rheological strength, with TriaC-18 exhibiting the lowest value (0.06), signifying superior gel strength. Gelation was observed only in hydrogelators with hydrophobic tails of at least 12 carbon atoms, with gel strength increasing with tail length. Notably, TriaC-18 hydrogel demonstrated exceptional adhesion properties, adhering well to surfaces (Figure 1F, G), suggesting potential for topical applications. The TriaC-18 hydrogel also exhibited prolonged stability, retaining favorable viscoelastic properties, including a tan δ value of 0.08, after 30 days of storage under ambient conditions with light exposure and in airtight enclosures (Figure S15). Evaluations of compressibility and injectability highlighted its practical utility, with no phase separation observed upon extrusion from plastic tubes or injection syringes (Figure 1H, I). These properties suggest that TriaC-18 is well-suited for both topical and localized therapeutic applications.

**Figure 1.**
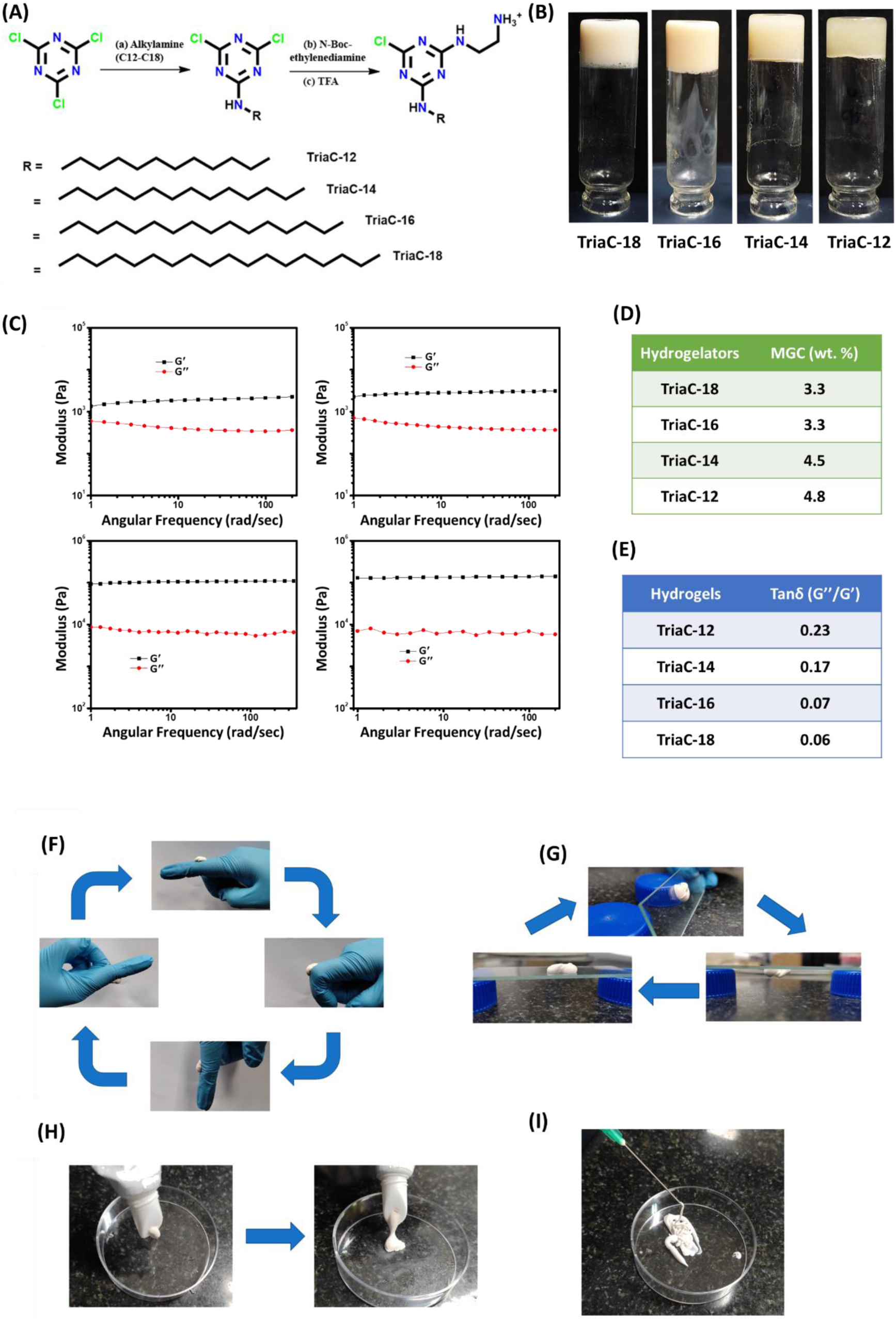
Synthesis, formation, and characterization of hydrogelators. (A) Synthetic pathway illustrating the synthesis of hydrogelators TriaC-18, TriaC-16, TriaC-14, and TriaC-12. (B) Photograph of inverted vials demonstrating successful hydrogel formation. (C) Dynamic rheological analysis via frequency sweep to determine the viscoelastic properties of the hydrogels. (D, E) Data on gelation, including minimum gelation concentrations. (F, G) Evaluation of adhesion properties specific to TriaC-18, (H, I) Characterization of the extrusion and injectability properties of TriaC-18, demonstrating its potential for use in applications.

### Morphological study

SEM and TEM analyses were performed to evaluate the morphology of the self-assembled aggregates. The results revealed fibrous, highly cross-linked structures, corroborating the formation of hydrogels (Figure 2A, B).

**Figure 2.**
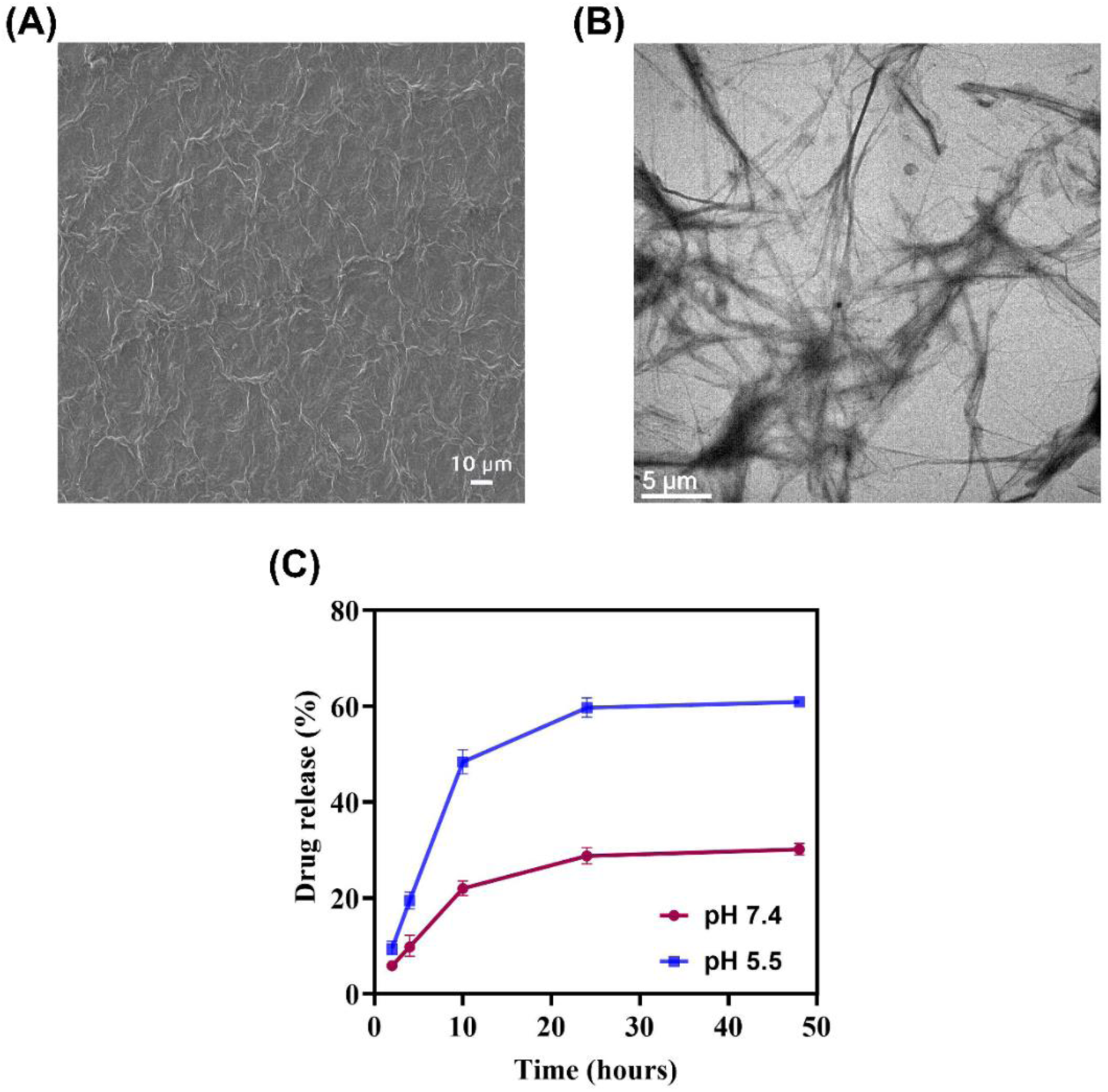
(A) SEM image of TriaC-18. (B) TEM image of TriaC-18. (C) pH-dependent DOX release study.

### DOX loading and controlled release

To evaluate the drug loading and release capabilities of the synthesized hydrogelator TriaC-18, doxorubicin, a widely utilized chemotherapeutic agent, was selected. Doxorubicin was successfully encapsulated within the TriaC-18 hydrogel, achieving a loading capacity of 73% after 96 hours (Figure S16). The drug release profile was then assessed under acidic (pH 5.5, mimicking the tumor microenvironment) and physiological (pH 7.4) conditions over time intervals ranging from 2 to 48 hours. Calibration curves established at pH 5.5 and pH 7.4 were used to quantify drug release. As shown in Figure 2C, drug release at mildly acidic pH was significantly higher compared to neutral pH. After 48 hours, 60.4% of the drug was released at pH 5.5, while only 27.7% was released at pH 7.4. These findings suggest that in the mildly acidic environment of the tumour microenvironment (TME), the gel can disassemble, facilitating a greater release of drug molecules compared to that observed under normal physiological pH conditions (pH 7.4), where the gel assembly remains more compact.

### Biological Studies

#### Dox encapsulated hydrogel matrix augments its anticancer activity against melanoma cell lines

Following the successful synthesis of hydrogelators and the characterization of their hydrogelation, drug loading, and release properties, further studies were performed using TriaC-18. The cellular cytotoxicity of doxorubicin (DOX), TriaC-18, and DOX-loaded TriaC-18 was assessed using MTT assays. Notably, DOX-loaded TriaC-18 exhibited a synergistic cytotoxic effect in both A375 (human melanoma) and B16-F10 (murine melanoma) cell lines. TriaC-18 alone displayed moderate cytotoxicity, with IC50 values of 10.7 µg/mL and 12.4 µg/mL in A375 and B16-F10 cells, respectively, which were higher than those observed for free DOX (IC50 = 1.4 µg/mL and 1.8 µg/mL in A375 and B16-F10 cells, respectively) (Figure 3A-B). Interestingly, the cytotoxicity in noncancerous HEK293 cells was markedly lower, suggesting that TriaC-18 demonstrates selective toxicity toward cancer cells. Enhanced, time-dependent cellular mortality was observed when doxorubicin was conjugated with TriaC-18 over a 48-hour period (Figure 3C(i), (ii)). Furthermore, scratch assays performed at four time points (0, 18 and 24 hours) revealed a significantly larger cell-free area in cultures treated with DOX-loaded TriaC-18 compared to those treated with free doxorubicin (1.4 µg/mL). Furthermore, treatment with 10.7µg/ml TriaC-18 demonstrated the capacity to suppress cellular migration, albeit to a lesser extent than doxorubicin alone (Figure 3D). These findings suggest that employing TriaC-18 as a carrier for doxorubicin may augment its inhibitory effects on cell migration, presenting a promising strategy for developing advanced cancer therapeutics aimed at mitigating metastasis. Additionally, a synergistic effect was observed between 1.4 µg/mL doxorubicin and 10.7 µg/mL TriaC-18 in a colony formation assay. Significant differences in colony formation were noted among the DOX-treated, TriaC-18-treated, and control groups. Remarkably, the application of DOX-loaded TriaC-18 led to complete inhibition of cell proliferation, highlighting the potent inhibitory potential of this combination therapy (Figure 3E).

**Figure 3.**
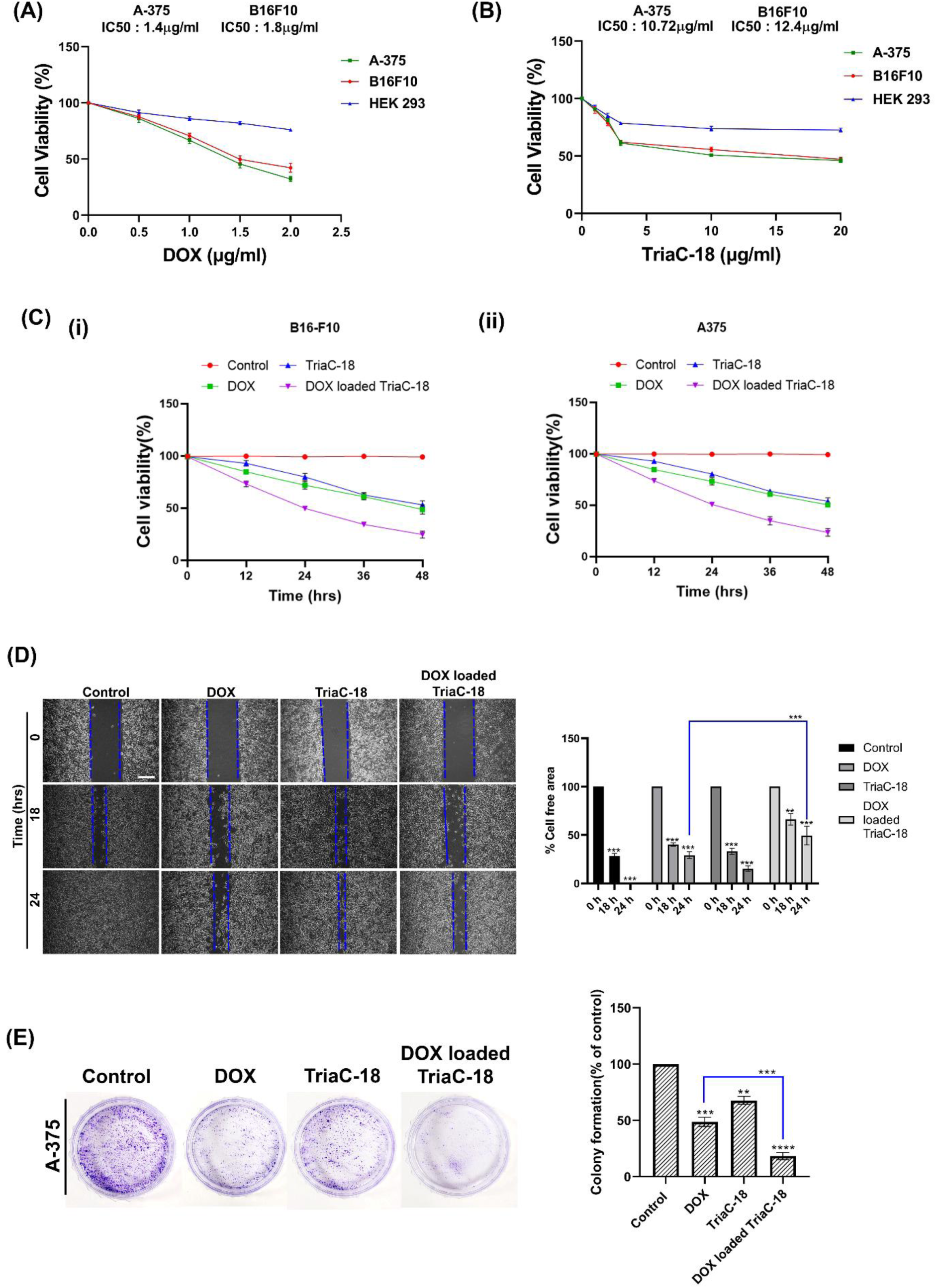
Anticancer activity of TriaC-18 & DOX loaded TriaC-18 on human melanoma cell line (A375) & murine melanoma cell line (B16-F10). (A) MTT assay of the DOX in different concentrations on A375, B16-F10 & HEK 293 cell lines. (B) MTT assay of the different concentrations of TriaC-18 on A375, B16-F10 & HEK 293 cell lines. (C) MTT assay of the DOX-loaded TriaC-18 on A375 & B16-F10 cells. (D) Scratch assay or cell migration assay at 0, 18 & 24 hrs. time point in the presence of 1.4 µg/ml DOX, 10.7 µg/ml of TriaC-18, and DOX loaded TriaC-18 in A375 cells and the percentage of cell-free area calculated. Scale bar = 20 µm (E) Colony formation assay of A375 cells in the presence of 1.4 µg/ml DOX, 10.7 µg/ml of TriaC-18, and DOX-loaded TriaC-18 and the percentage of colony is calculated. Bar graphs are represented as mean +/− SD (n=3). Where *<0.05, **<0.01, ***<0.001, ****<0.0001 & ns is non-significant.

In conclusion, this study highlights the potential of TriaC-18 as an effective carrier for enhancing the therapeutic efficacy of doxorubicin in melanoma treatment. The observed synergistic effects of DOX-loaded TriaC-18 in suppressing cell proliferation and migration underscore its promise as a potent strategy for targeting metastasis. Moreover, the reduced cytotoxicity of TriaC-18 in noncancerous cells indicates a favorable safety profile.

#### Nuclear localization facilitates precise & efficient delivery into the nucleus

Doxorubicin possesses an intrinsic ability to translocate into the nucleus and induce double-strand DNA breaks, ultimately leading to apoptosis. However, its nuclear translocation efficiency is relatively low. Our previous findings demonstrated that doxorubicin-loaded TriaC-18 hydrogelator significantly increased cell death, suggesting that the hydrogelator may facilitate more efficient cellular uptake of doxorubicin. To investigate this possibility, A375 cell lines were treated with free doxorubicin (1.4 µg/mL) and doxorubicin-loaded TriaC-18 for 2 and 4 hours. Doxorubicin showed red fluorescence so we treated the cells with 1.4 µg/ml free doxorubicin and doxorubicin-loaded TriaC-18 along with Hoechst 33342, a cell permeable nuclear dye. Using fluorescence microscopy imaging we found that free doxorubicin took 4 hrs to translocate into the nucleus but when doxorubicin loaded TriaC-18 was added in the cell culture medium, doxorubicin translocated into the nucleus within 2 hrs (Figure 4A). Nuclear condensation is a characteristic phenomenon of cells undergoing apoptosis. Thus, to probe programmed cell death we further performed a nuclear condensation assay using fluorescence microscopy imaging, we stained the cells with Hoechst 33342 after 24 hrs incubation with doxorubicin and TriaC-18 loaded doxorubicin, Figure 4B clearly shows that doxorubicin treatment sequestered DNA in single nuclear bodies which depicted the condensation of the DNA molecules, the amount of nuclear condensation increases in the cells which are treated with DOX loaded TriaC-18.

**Figure 4.**
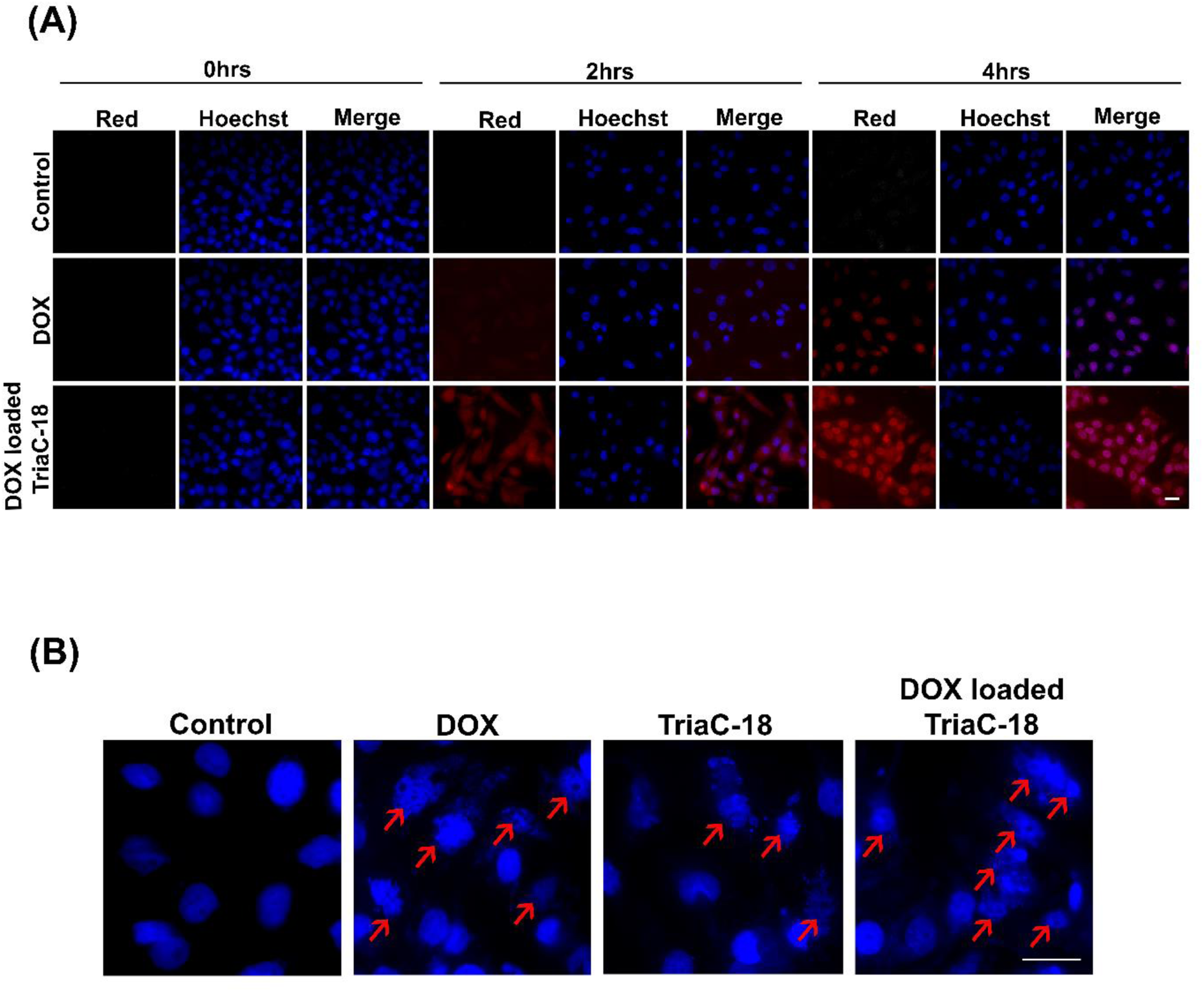
Cellular uptake of free DOX and DOX loaded in TriaC-18 hydrogelator. (A) Fluorescence microscopy imaging of DOX uptake and translocation into the nucleus in different time points (0, 2 and 4 hrs.) Scale bar = 20 µm (B) Nuclear condensation assay of A375 cells after treating with 1.4µg/ml DOX, 10.7 µg/ml TriaC-18 and DOX loaded TriaC-18. Scale bar = 20 µm.

#### DOX-loaded TriaC-18 substantially augments the DOX-induced Apoptosis

Based on the preceding data, we have established the anti-cancer properties of doxorubicin-loaded TriaC-18, free doxorubicin and free TriaC-18. An effective anticancer agent is required to impede cell division, thereby arresting the unrestrained and rapid proliferation of cancerous cells. We tried to investigate further to gain a comprehensive understanding of which specific phase of the cell cycle was targeted by doxorubicin-loaded TriaC-18, free doxorubicin, and free TriaC-18. As deduced from our extensive literature review, it is well-established that doxorubicin possesses the capability to induce cell cycle arrest at the G2/M phase, concomitant with inducing cellular apoptosis. To elucidate the effects of our experimental conditions, we conducted a cell cycle analysis following a 24-hour treatment of A375 cells with 1.4 µg/ml doxorubicin, 10.7 µg/ml TriaC-18, and doxorubicin-loaded TriaC-18. As depicted in Figure 5A, doxorubicin exhibited its characteristic ability to arrest the cell cycle at the G2/M phase. However, under the other treatment conditions, cell cycle arrest was not observed. Interestingly, in the case of cells treated with doxorubicin-loaded TriaC-18, there was a notable increase in the Sub G0 population, indicating a synergistic effect resulting in enhanced cellular apoptosis. Furthermore, in cells treated with free TriaC-18, a similar increase in the Sub G0 population was observed, suggesting that TriaC-18 alone possesses some capacity to induce cell death independently. Treatment with 1.4 µg/ml doxorubicin, 10.7 µg/ml TriaC-18, and doxorubicin-loaded TriaC-18 led to a notable increase in the Sub G0 cell population, unequivocally signifying their potent induction of apoptosis. As a hallmark of apoptotic events, the translocation of phosphatidylserine (PS) from the inner to the outer leaflet of the cell membrane was discerned through FITC-tagged Annexin V binding to exposed PS residues, while propidium iodide (PI) translocated into the nucleus to stain DNA molecules, indicative of increased membrane permeability in late apoptosis stages. To provide further validation of their apoptotic efficacy, we conducted Annexin V-PI assays using Flow cytometry. While 24 hrs of 1.4µg/ml doxorubicin treatment elevated the Annexin V and PI positive cell population by approximately 12.1% and 10.7µg/ml TriaC-18 treatment increased it by approximately 6.4%, doxorubicin-loaded TriaC-18 treatment synergistically enhanced the apoptotic cell population, with approximately 22.6% of cells exhibiting Annexin V and PI positivity (Figure 5B).

**Figure 5.**
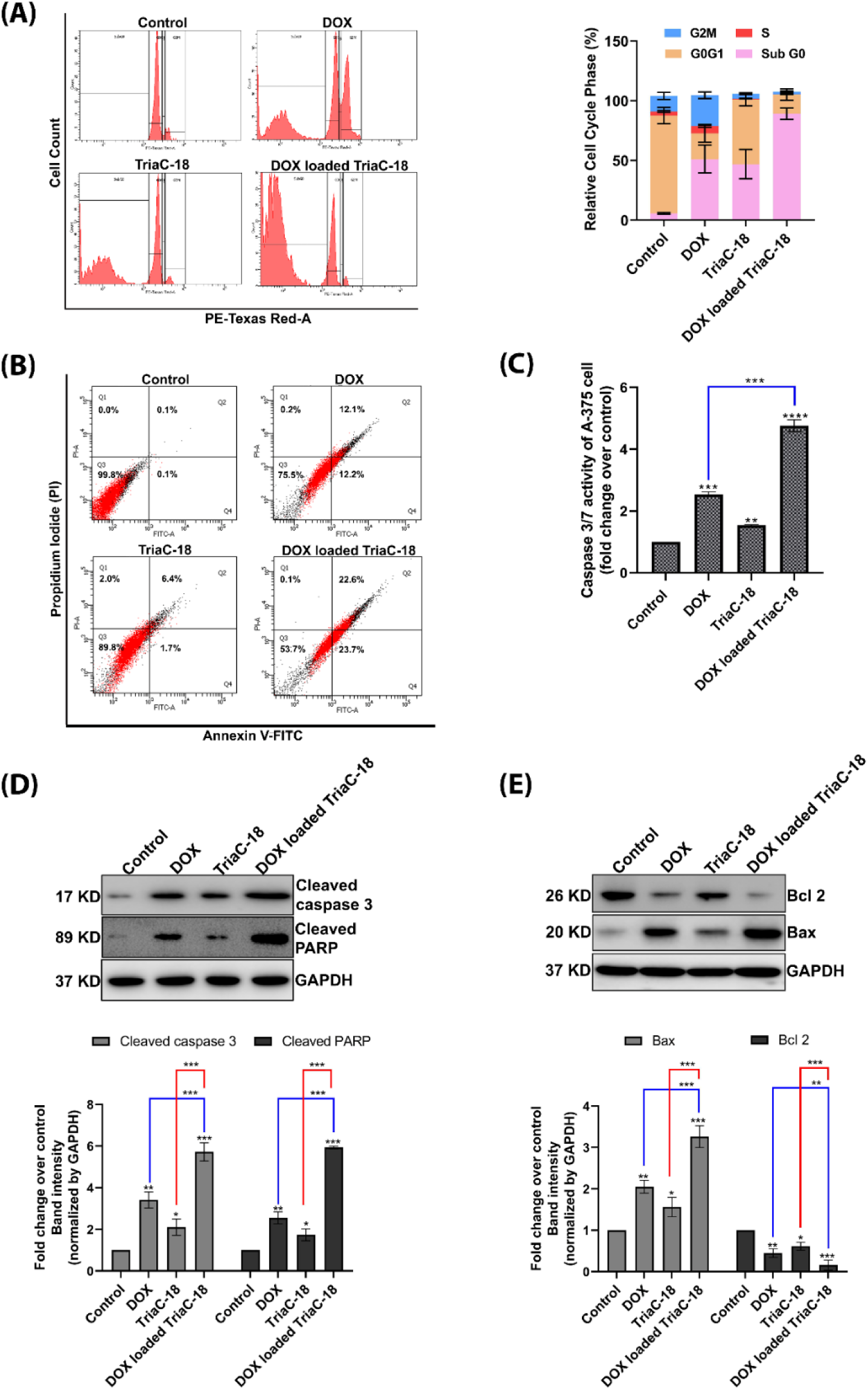
Apoptotic potential of 1.4 µg/ml DOX, 10.7 µg/ml TriaC-18 & DOX loaded TriaC-18 in A375 cells. (A) Flowcytometric Cell cycle assay using propidium iodide (PI) in A375 cells. (B) Annexin-V and PI assay using flowcytometry of A375 cells. (C) Caspase3/7 colorimetric assay of A375 cells. (D) Western blot analysis of apoptosis marker cleaved caspase3 and cleaved PARP in A375 cells and densitometric analysis represented as bar diagram calculated using ImageJ and GraphPad Prism 8. (E) Western blot analysis of antiapoptotic protein Bcl2 and proapoptotic protein Bax of A375 cells. Bar graphs are represented as mean +/− SD (n=3). Where *<0.05, **<0.01, ***<0.001, ****<0.0001 & ns is non-significant.

Apoptosis, a well-established cellular process, is typically initiated by the activation of effector Caspases, such as Caspase 3/7, ultimately leading to the degradation of critical cellular components, including DNA, Golgi bodies, and the nucleus cytoskeleton. To explore this further, we assessed Caspase 3/7 activity in 1.4 µg/ml doxorubicin, 10.7 µg/ml TriaC-18, and doxorubicin-loaded TriaC-18-treated A375 cells. As cells underwent apoptosis, Caspase 3/7 activation ensued, processing the Acetyl-Asp-Glu-Val-Asp p-nitroanilide (Ac-DEVD-p-NA) substrate, which absorbed light at 360 nm wavelength when apoptosis occurred. Remarkably, doxorubicin-loaded TriaC-18 markedly amplified Caspase 3/7 activity compared to free doxorubicin, while intriguingly, the hydrogelator TriaC-18 also induced a slight increase in Caspase 3/7 activity (Figure 5C). These findings highlight the robust pro-apoptotic effects of doxorubicin-loaded TriaC-18, making it a promising candidate for cancer therapy. Then we went on to check the cleaved Caspase 3 and cleaved PARP, the marker of apoptosis, the expression pattern in doxorubicin, TriaC-18, and doxorubicin-loaded TriaC-18 treated A375 cells. We also checked the expression pattern of the anti-apoptotic protein Bcl2 and pro-apoptotic protein Bax in above mentioned treatment conditions. We found that the doxorubicin-loaded TriaC-18 treatment increased the cleaved Caspase3 and cleaved PARP expression (Figure 5D). There was a dramatic increase and decrease in the expression of the pro-apoptotic protein Bax and the anti-apoptotic protein Bcl2, respectively (Figure 5E). These findings highlight the potent pro-apoptotic effects of doxorubicin-loaded TriaC-18, underscoring its potential as a promising approach for advancing cancer therapeutic strategies. Interestingly, TriaC-18 also exhibited inherent pro-apoptotic activity, prompting further investigation into the underlying mechanisms and key molecular targets driving TriaC-18-induced cell death in the A375 melanoma cell line.

#### Molecular docking indicates potential interaction between TriaC-18 and PI3K catalytic subunit p110α

In the above findings, we observed that triazine-based hydrogelator TriaC-18 not only enhances the efficiency of DOX by facilitating its cellular and nuclear transport but also seems to induce programmed cell death itself. Several 1,3,5-triazine-containing compounds have demonstrated potent inhibition of PI3K/mTOR and are undergoing various stages of drug development^24^. Notable examples include Gedatolisib (PKI-587), a highly potent dual inhibitor of PI3Kα/γ, and mTOR^54^. Another triazine-based drug, Buparlisib (BKM120) inhibits all the PI3K isoforms p110α, β, δ, and γ^55^ ^56^. Some of the triazine-based drugs are shown in Scheme 2. To investigate the mechanism underlying the impact of our synthesized hydrogelator TriaC-18 on cancer progression, we performed a molecular docking simulation to check if the compound could have some interaction with PI3K, and therefore potentially inhibit the PI3K pathway which is considered among the most common dysregulated pathways in most cancers.

Molecular docking using AutoDock 4.2 indicated a moderately strong binding affinity of −6.34 kcal/mol with the PI3K catalytic subunit p110α. Although the compound did not form any hydrogen bond with the hinge region Val851, which is a characteristic of several kinase inhibitors^57^, the TriaC-18 alkyl tail seemed to be involved in a significant number of hydrophobic interactions with several residues including Trp780, Ile800, Ile848, Val850, Phe930, Ile932, Asp933 as well as the hinge region Val851 (Fig 6B, C). Apart from this, hydrogen bonds were formed with Lys802, Asp805, Asp810, and the DFG loop residues Asp933 and Phe934, and a π-anion interaction also seemed to take place between the triazine ring and Asp805 (Fig 6B, C). This DFG loop binding is one of the chief interactions that stabilize the binding of an ATP molecule in the active site^58^. If an inhibitor can form a stable interaction with it, it could potentially compete with ATP in binding to the active site. This provides a preliminary insight into a possible molecular interaction that could take place between TriaC-18 and the active site of PI3K, through which the Compound could exhibit its pro-apoptotic effect on cancer cells.

**Figure 6.**
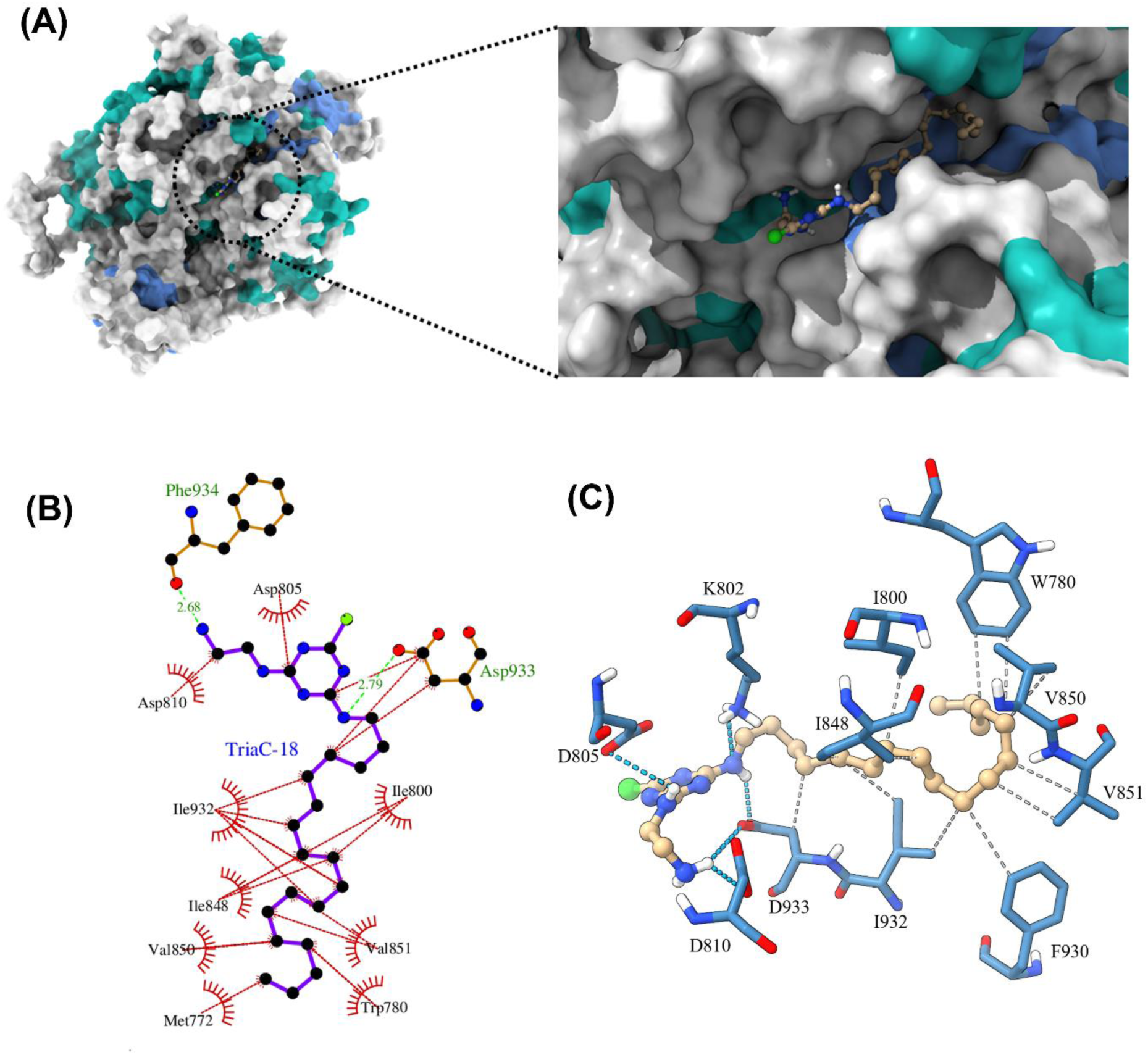
Molecular docking of TriaC-18 in PI3K (A) TriaC-18 docked in the active site of PI3K catalytic subunit p110α (PDB ID: 6OAC) (B) 2-D interaction profile of the ligand TriaC-18 with the active site of the protein (C) 3-D interaction profile generated using PLIP (Protein-Ligand Interaction Profiler); blue – hydrogen bonding, grey dashed – hydrophobic interactions.

**Scheme 2.**
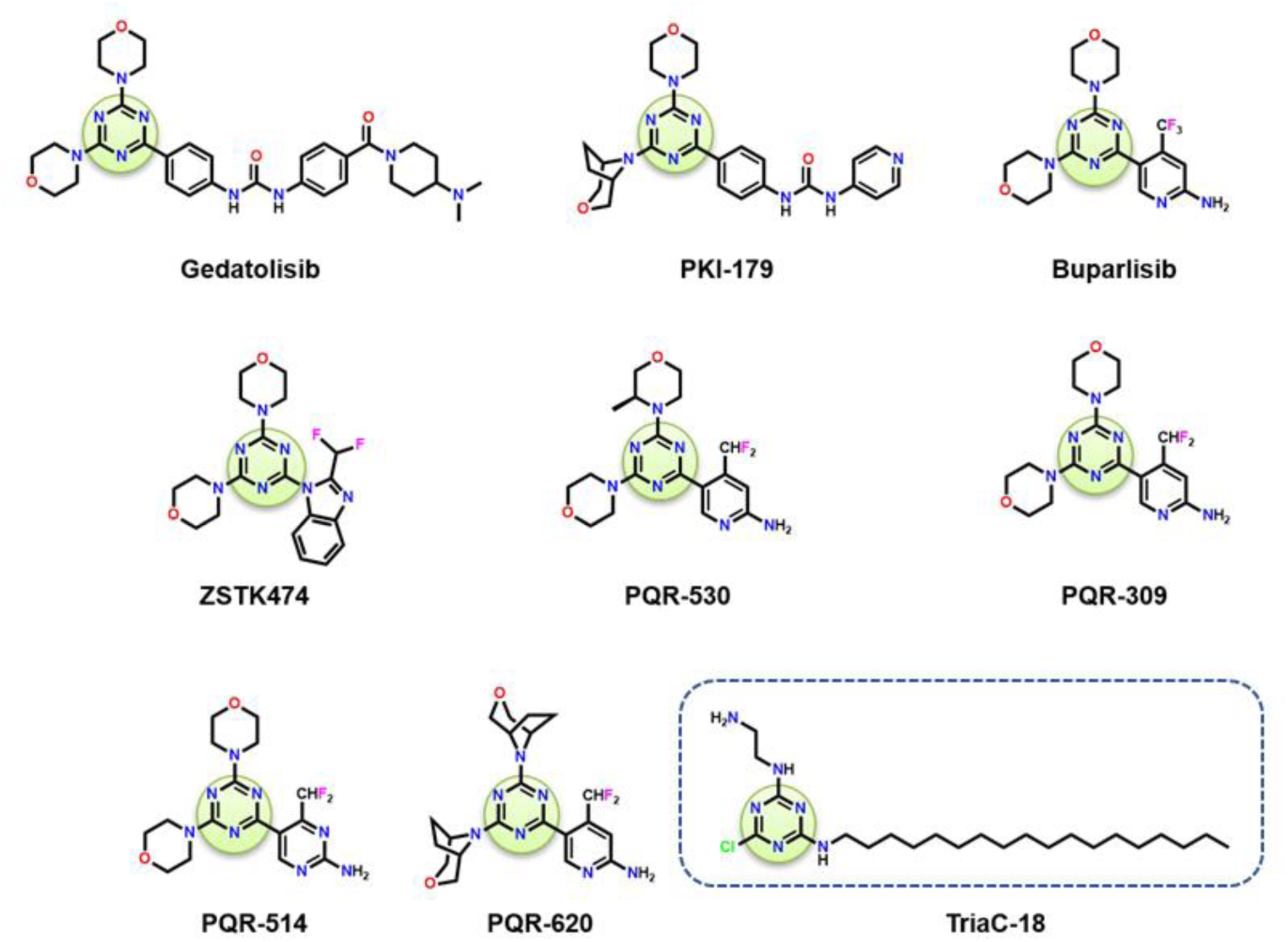
Chemical structures of some triazine-based PI3K inhibitors and the hydrogelator TriaC-18.

#### Molecular dynamics simulation studies show stable binding of TriaC-18 in the active site of PI3K

The kinase family of enzymes in general has a hinge region that has a hydrophobic pocket where the adenosine group of ATP binds, and a hydrophilic pocket region, where the triphosphate tail sits. As ATP anchors to the kinase hinge region, the purine ring forms hydrogen bonds with the backbone of the hinge region, and the triphosphate group is stabilized by a stretch of Asp-Phe-Gly sequence, known as the DFG motif^59^. In general, kinase inhibitors that bind to the active site of a kinase need to bind strong enough so that nanomolar concentrations of the drug molecule can competitively inhibit the binding of millimolar concentrations of ATP inside the cell. As a result, it is important for a competitive inhibitor to form stronger and possibly even additional interactions with the active site than that formed by ATP. Most kinase inhibitors, including the triazine-based PI3K inhibitors, involve hydrogen bonding interactions with the hinge region Val851 residue as well as the DFG loop and form stable interactions^60^ which inhibit the binding of ATP. It was thus important to probe into the interactions and dynamics of the hydrogelator TriaC-18 docked in the active site of PI3K catalytic subunit p110α.

To validate the molecular docking result of the compound TriaC-18 in the active site of PI3K catalytic subunit p110α and to gain more insights into the possible dynamics and interactions of the compound with the active site residues, a comparative molecular dynamics simulation study between the docked complex containing TriaC-18 and the protein without the docked compound (APO) as a control was performed.

#### Root mean square deviation (RMSD)

To understand the structural changes of the PI3K catalytic subunit p110α in ligand (TriaC-18) bound and unbound conditions, we computed the RMSD of the protein backbone with respect to its energy-minimized conformation. Figures 7B and C show RMSD of the protein backbone at temperature 310 K in the absence and the presence of TriaC-18 respectively. RMSD values show that the protein’s structure is stabilized throughout the simulation run (250 ns). Furthermore, we calculated the average RMSD (RMSD_avg_) from the last 100ns of the simulation run. The RMSD and RMSD_avg_ are reported in Supplementary Table 2 and Figure S13A of supplementary information. The difference in this average RMSD value for the two systems was calculated to be about 0.456 Å.

**Figure 7.**
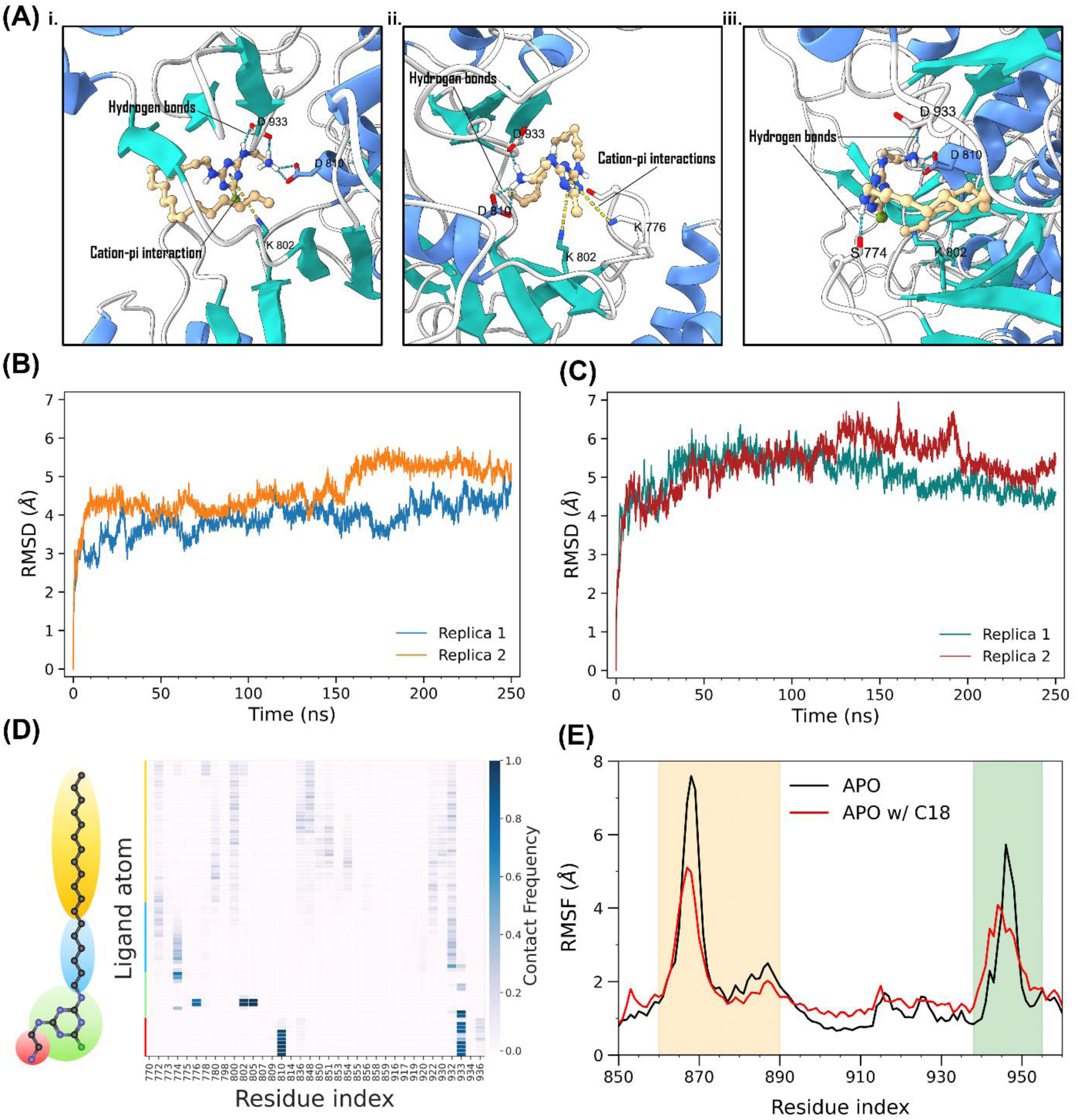
Molecular dynamics simulation results of the TriaC-18 (C18) docked in the active site of PI3K catalytic subunit p110α (A) Cartoon representation of the interactions between the active site residues and TriaC-18: Cation-pi interaction (yellow dashed line) of the triazine ring of TriaC-18 and residues K776 and K802 (i and ii), hydrogen bonding interaction (blue dashed line) with S774, D810 and D933 (i, ii and iii). (Atom color codes: Tan – C, Blue – N, Red – O, Green – Cl, White – H) (B) RMSD profile of two replicates of control (APO, without the inhibitor) for 250 ns (C) RMSD profile of two replicates of the inhibitor (TriaC-18) bound complex for 250 ns (D) Contact frequency heatmap showing the frequency of contact of all the active site residues (within 4 Å of the ligand) with the different atoms of the ligand. The different parts of the ligand have been highlighted in different colors to highlight the corresponding regions in the heatmap. (E) RMSF profile of the TriaC-18 bound complex (APO w/ C18), along with the control (APO), focused on the residues 850 to 960.

#### Root mean square fluctuation (RMSF)

For a detailed understanding of the structural changes in the proteins, we have performed RMSF calculation of Cα atoms of all the residues. Figure S13B shows the RMSF of Cα atoms at temperature 310 K in the ligand (TriaC-18) bound and unbound conditions. However, we did not observe any significant changes in the RMSF values except for a few specific residues. Highlighting the residues near the catalytic site that show significant changes in the ligand-bound condition, Figure 7E focuses on the two regions, i.e., residues 860 to 890 and residues 942 to 955. The binding of the ligand TriaC-18 leads to increased stability of these two regions. However, these residues are not in direct contact with the ligand, indicating the allosteric effect of ligand binding. We can also relate this observation with previous findings, where it was shown that the substrate binding leads to a decrease in the fluctuation of residue around the active site tunnel^61^.

#### Contact frequency of the residues

In order to understand the interaction between the compound TriaC-18 and the active site residues of the protein, we calculated the contact frequencies of each of the residues in contact (within 4 Å) with the different atoms of TriaC-18. The contact frequency heatmap (Figure 7D) showed some stable interactions that persisted for almost the entire simulation time. One of them was the interaction between the terminal amine and the Asp933 and Asp810 residues, which suggests stable hydrogen bonding and salt-bridge formation (Figure 7A). Stable contacts were also observed between the triazine nitrogen atoms of TriaC-18 and the Lys802 and Lys776 residues, which indicates cation-pi interactions of the protonated amine of the lysine residues and the triazine group. A less frequent, but moderately stable interaction was observed between the N1 atom (secondary amine nitrogen atom linking the triazine group and the 18-carbon tail) of TriaC-18 and the Ser774 residue, which suggests stable hydrogen bonding. Apart from these polar interactions, there were also numerous, though less frequent, hydrophobic interactions between the 18-carbon tail of TriaC-18 and many hydrophobic residues of the active site, most notably with Arg770, Trp780, Ile800, Tyr836, Ile848, Val851 (hinge region valine residue), Met922, Phe930, and Ile932. Additionally, hydrogen bond distribution analysis showed the formation of four to five hydrogen bonds, among which those with the residues Asp810 and Asp933 were the most stable, and those with the residue Ser774 remained moderately stable (Figure S13B). In summary, the molecule seemed to attain a stable orientation with the 18-carbon alkyl chain oriented towards the hinge region hydrophobic pocket where it engaged in numerous hydrophobic interactions while the triazine ring along with the ethylene diamine moiety oriented towards the hydrophilic pocket where it interacted with the DFG loop via hydrogen bond and possible salt bridge formation, which could potentially inhibit ATP binding.

#### Incorporation of Dox into the hydrogel matrix unleashes its anticancer potential, enhancing chemotherapy efficacy even at reduced doses

Molecular docking and molecular dynamics simulation studies demonstrated a stable interaction of TriaC-18 with the active site of PI3K, a kinase responsible for converting PIP2 into PIP3 within the cell membrane. Subsequently, PIP3 activates the downstream signaling molecule AKT, which leads to the phosphorylation of the latter. Phosphorylated AKT (p-AKT) in turn, leads to the inactivation of FOXO-3a via phosphorylation, followed by proteasomal degradation. Inhibition of PI3K results in the translocation of unphosphorylated FOXO-3a into the nucleus, where it orchestrates the specific transcriptional synthesis of BH3 family pro-apoptotic proteins, including Bim, PUMA, and NOXA. Notably, DOX primarily translocated into the nucleus and induces double-stranded DNA breaks, consequently elevating p53 expression and activating PUMA/NOXA, the sole BH3 family proapoptotic protein. To validate these, we treated A375 cells with 10.7µg/ml TriaC-18 and 1.4µg/ml DOX and monitored AKT phosphorylation over time (0-24 hrs) via Western blot analysis. Our results revealed that TriaC-18 inhibited AKT phosphorylation after 6 hrs, while DOX required 12 hrs for this effect (Figure 8A, B, Figure S14 A, B). Additionally, we assessed AKT and FOXO-3a activation and inactivation in cells treated with DOX, TriaC-18, and DOX-loaded TriaC-18 (Figure 8C). Remarkably, the 10.7 µg/ml TriaC-18 exhibited the superior capacity to inhibit AKT activation, induce FOXO-3a inactivation via phosphorylation, and increase Bim expression, particularly pronounced in DOX-loaded TriaC-18-treated cells. While DOX induces dsDNA breaks, as evident from the early increase in γ-H2AX expression (Figure 8D, E, Figure S13 C), 10.7 µg/ml TriaC-18 alone did not induce dsDNA breaks. However, DOX-loaded TriaC-18 increased the effects of free DOX on dsDNA breaks and elevated PUMA/NOXA expression as TriaC-18 delivered a greater quantity of DOX into the cells (Figure 8F). Consequently, our findings suggest that TriaC-18 not only serves as an efficient DOX delivery agent in melanoma chemotherapy but also possesses intrinsic anticancer potential by inhibiting PI3K, contributing to the synergistic effects observed when using DOX-loaded TriaC-18 as a local chemotherapeutic agent in melanoma cell lines.

**Figure 8.**
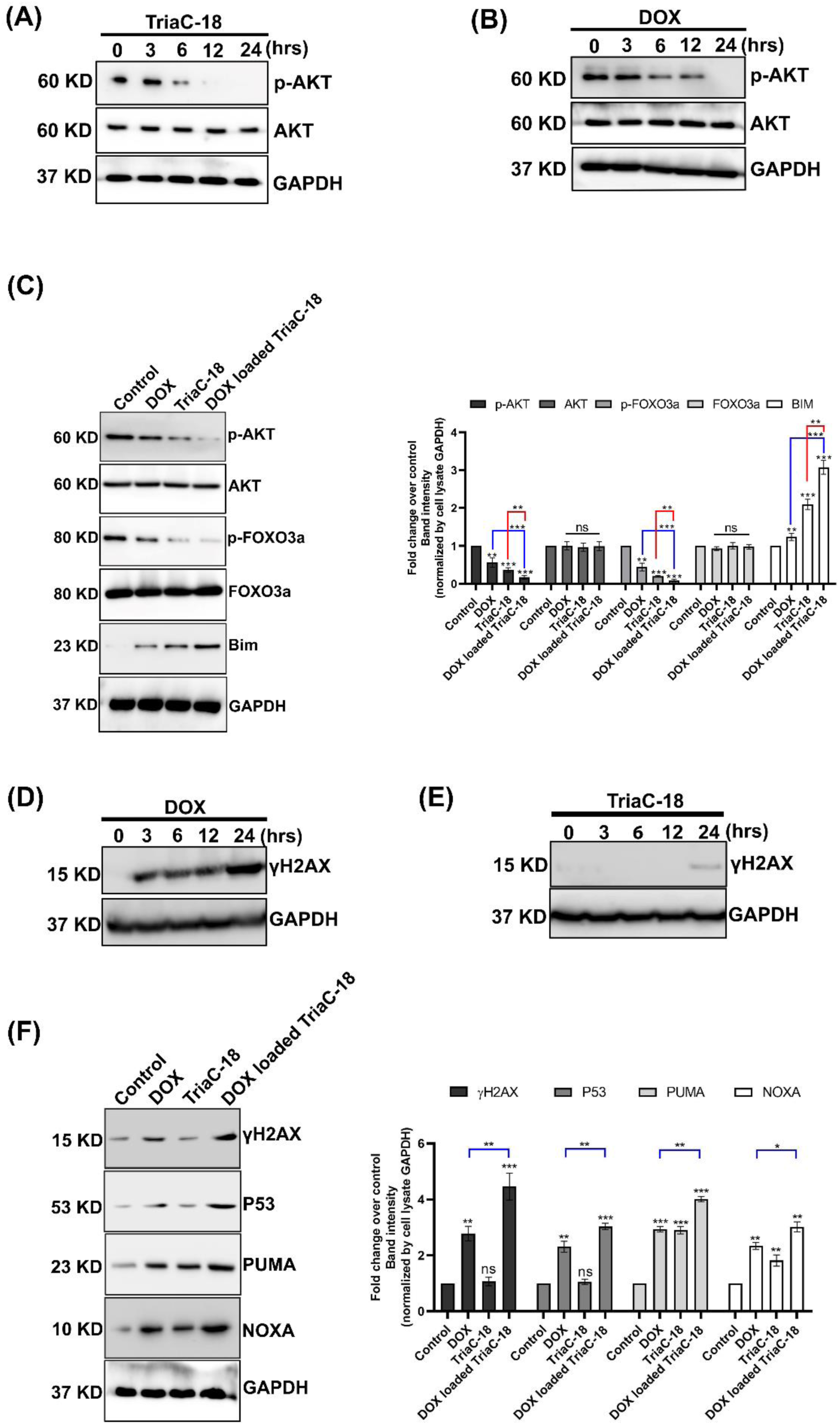
Molecular mechanism analysis of 1.4 µg/ml DOX, 10.7 µg/ml TriaC-18, and DOX loaded TriaC-18 mediated apoptosis. (A) Western blot analysis of AKT phosphorylation along with total AKT expression in different time points (0-24 hrs.) after treating A375 cells with 10.7 µg/ml TriaC-18. (B) Western blot analysis of AKT phosphorylation along with total AKT expression in different time points (0-24 hrs.) after treating A375 cells with 1.4 µg/ml DOX. (C) Western blot analysis of p-AKT, p-FOXO-3a, and Bim of A375 cells after treatment with 1.4 µg/ml DOX, 10.7 µg/ml TriaC-18, and DOX loaded TriaC-18. (D, E) dsDNA break marker γ-H2AX expression analysis using western blot of A375 cells after 1.4 µg/ml DOX and 10.7 µg/ml TriaC-18 treatment respectively in different time points (0-24 hrs). (F) γ-H2AX, P53, PUMA, and NOXA expression analysis of A375 cells after treatment with 1.4 µg/ml DOX, 10.7 µg/ml TriaC-18 and DOX loaded TriaC-18 using Western blot. Bar graphs are represented as mean +/− SD (n=3). Where *<0.05, **<0.01, ***<0.001, ****<0.0001 & ns is non-significant.

### TriaC-18 enhanced the chemotherapeutic potential of DOX in *In Vivo* system

Building on our *in vitro* findings, we tried to investigate the *in vivo* chemotherapeutic impact of combining DOX with TriaC-18 hydrogelator in a murine model (C57BL/6) with subcutaneously implanted B16-F10 cells. Post 7-day tumor growth, 3.3 wt% TriaC-18, 2.5 mg/kg DOX, and DOX-loaded TriaC-18 were administered via injection (single dose during the 35 days), and the ensuing 35-day observation period revealed intriguing results. The schematic for the *in vivo* experiment is shown in Figure 9A. Figure 9B pinpoints the injection sites for TriaC-18, DOX, and DOX-loaded TriaC-18. Interestingly, tumor growth was impeded significantly more in the DOX-loaded TriaC-18 treated mice compared to that in the control as well as 2.5 mg/kg DOX (Figure 9C). Additionally, a substantial improvement in mouse survivability was observed in the DOX-loaded TriaC-18 cohort (Figure 9D). It is worth noting that throughout the 20-day experiment, we saw no discernible change in the mice’s body weight, indicating that the 3.3 wt% TriaC-18, 2.5 mg/kg DOX, and DOX loaded TriaC-18 application may not have been toxic (Figure 9E). After 20 days, the mice were sacrificed and the tumor volume and weight were analyzed. We found that both the volume and the weight decreased significantly in DOX-loaded TriaC-18 treated mice compared to control and 2.5 mg/kg DOX treated mice (Figure 9F, G). We also performed H & E staining of various organs derived from the different sets of mice to check the toxicity profile of TriaC-18. The data revealed no significant changes in the cellular morphology and tissue arrangement which confirms the non-toxic nature of TriaC-18 (Figure S18). We have also checked the expression of various pro-inflammatory markers and we observed significant reduction in the expression of IFN-γ and TNF-α (Figure S19) upon Dox loaded TriaC-18 treatment. This comprehensive study elucidates the potential therapeutic impact of DOX-loaded TriaC-18 *in vivo*, paving the way for advanced cancer treatment strategies.

**Figure 9.**
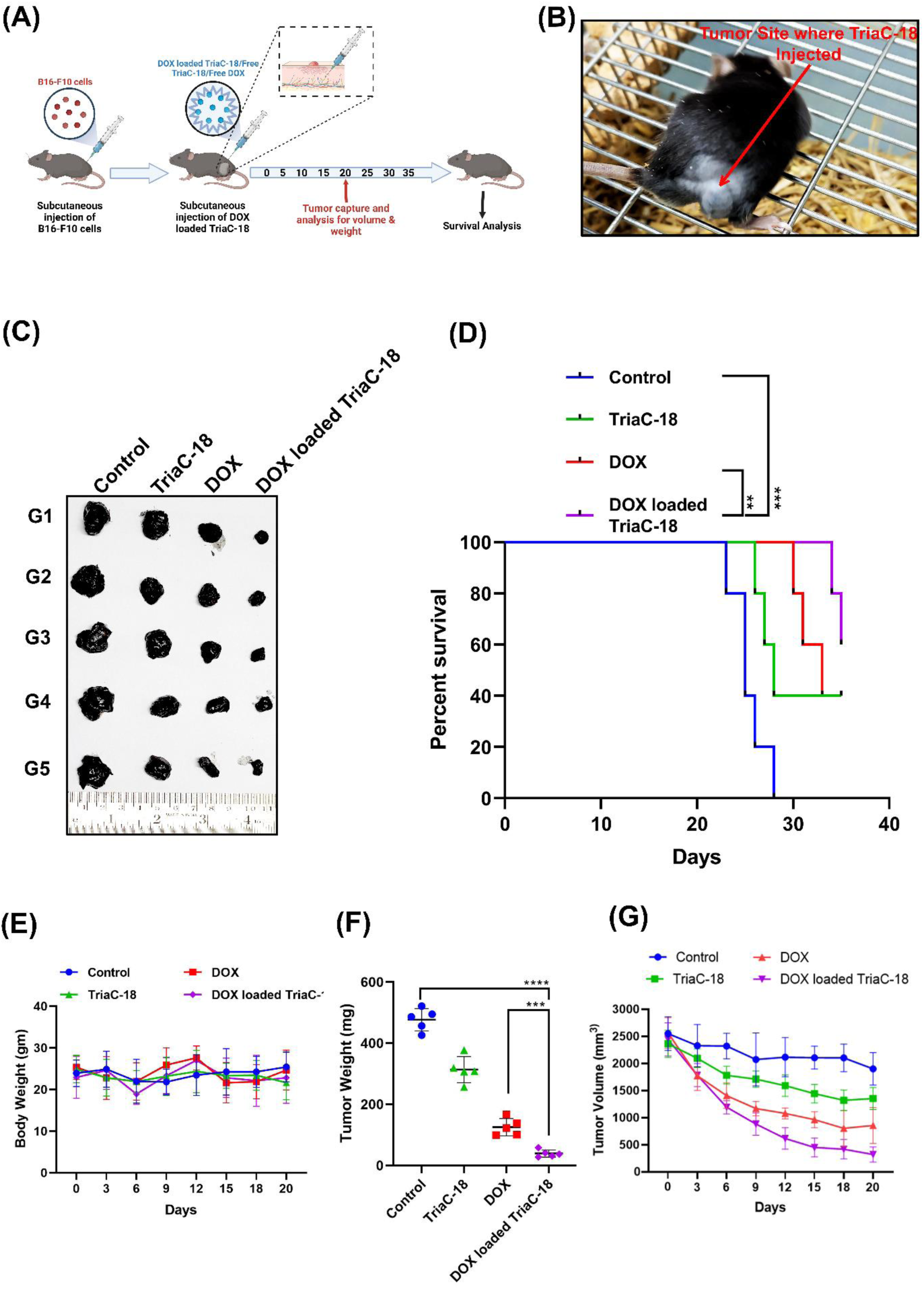
In vivo anticancer activity of TriaC-18 and DOX-loaded TriaC-18 in C57BL/6 mice (n=3). (A) Schematic representation of in vivo experiment. (B) The red arrow indicates the site of injection. (C) Representative tumor image in different treatment conditions. (D) Survival analysis of tumor-bearing mice during 35-day experiment period. (E) Body weight measurement of tumor-bearing mice treated with 2.5 mg/kg DOX, 3.3 wt% TriaC-18, and DOX-loaded TriaC-18 during the 35-day experiment period. (F-G) Volume and weight of the tumor measured after sacrificing the mice. Graphs are represented as mean +/− SD (n=5). Where *<0.05, **<0.01, ***<0.001, ****<0.0001 & ns is non-significant.

## CONCLUSION

1.3,5 Triazine is a biologically and medicinally active scaffold that is found in many FDA-approved drugs. In this work, we have designed and synthesized a series of triazine-based hydrogelators varying its carbon chain from C12-C18. Rheology data show that the gelation property of those hydrogelators decreases with a gradual decrease in the carbon chain. Highly cross-linked morphology obtained from SEM and TEM further confirms the gelation properties of those hydrogelators. We then used one of the hydrogelators, TriaC-18 as a potent delivery agent for DOX against A375 and B16-F10 melanoma cell lines. Subsequent confirmation studies, such as MTT assay, scratch assay, and colony formation assay, demonstrated that DOX-loaded TriaC-18 displayed remarkable potential for cancer cell eradication. Microscopic analysis further revealed that TriaC-18 enhanced the cellular transport of DOX, with translocation into the nucleus achieved within only 2 hours when encapsulated within the hydrogelator, TriaC-18. Cell cycle analysis, Annexin V-PI assay, Caspase 3/7 activity assay, and western blot analysis of cleaved Caspase3 and cleaved PARP collectively indicated that DOX-loaded TriaC-18 exhibited greater capacity to initiate apoptosis compared to free DOX. Notably, TriaC-18 also manifested a degree of inherent anticancer activity. Molecular docking and molecular dynamics studies revealed a potentially stable interaction between TriaC-18 and the active site of PI3K catalytic subunit p110α. Several stable interactions, including cation-pi interactions, hydrophobic interactions, and hydrogen bonds formed throughout the course of the molecular dynamics simulation which seemed to stabilize the TriaC-18 molecule in the active site. The interaction with and inhibition of PI3K was further evident from experimental findings. Western blot analysis conducted on a range of proteins, including AKT, FOXO-3a, and Bim, demonstrated notable variations in their expression levels following exposure to TriaC-18 treatment and treatment with DOX-loaded TriaC-18, which indicates that TriaC-18 was indeed having an inhibitory effect on the PI3K signaling pathway. In contrast, DOX-induced apoptosis through DNA damage. Therefore, our findings highlight the potential for designing and synthesizing triazine-based small molecules with hydrogelation properties to serve as efficient drug carriers. Moreover, these molecules exhibit intrinsic anticancer activity, as demonstrated by their cytotoxic effects in this study. Further exploration and optimization of such compounds could pave the way for the development of innovative drug delivery systems with enhanced anticancer efficacy, contributing to advancements in cancer therapeutics. In conclusion, the multifaceted approach adopted in this study highlights the potential of TriaC-18 as a versatile platform for the targeted delivery of doxorubicin in melanoma therapy. The combination of TriaC-18’s intrinsic anticancer properties and its ability to enhance the cellular transport of doxorubicin positions it as a promising candidate for further investigation in cancer treatment (Scheme 3).

**Scheme 3.**
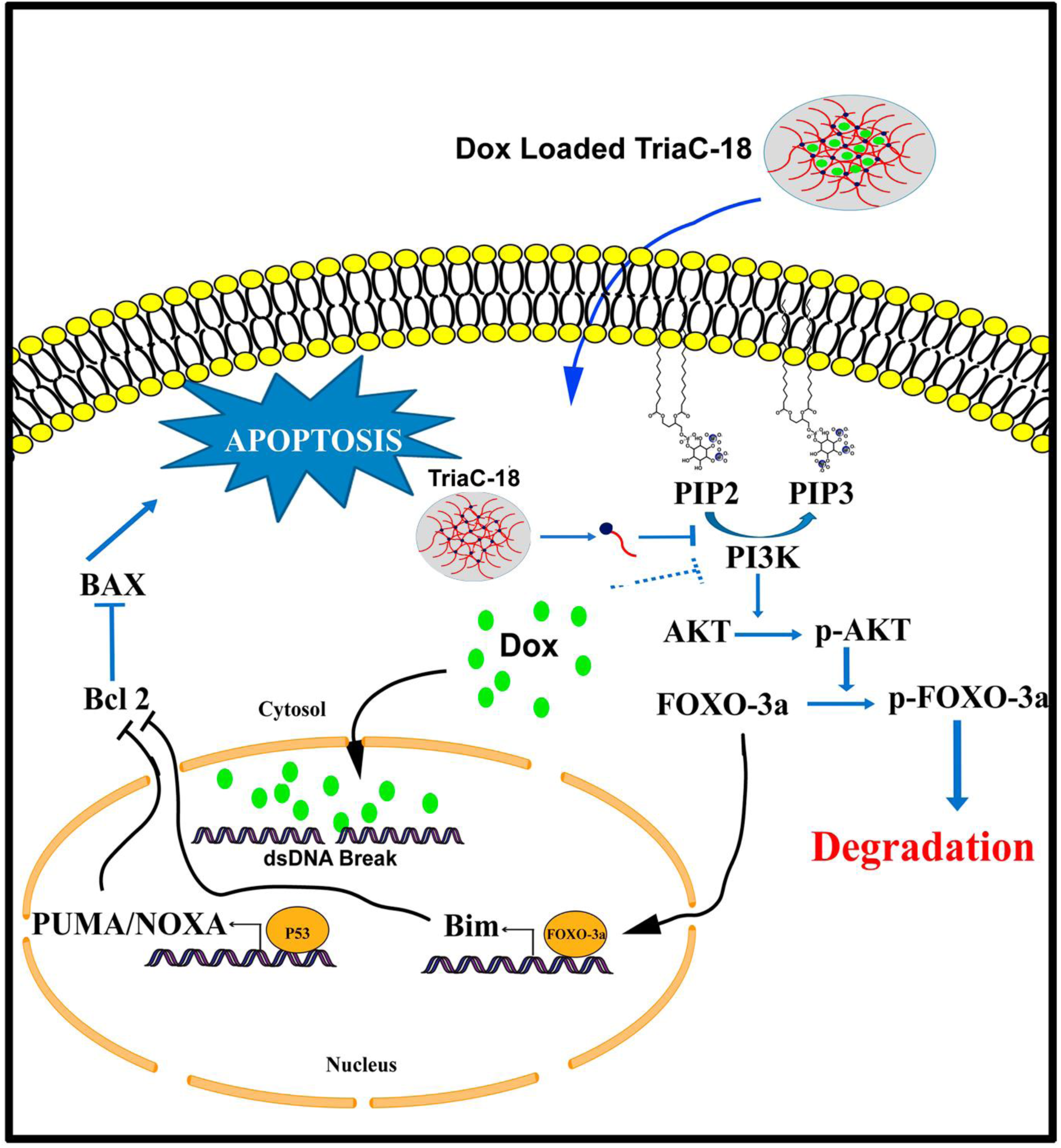
Mechanism of action of DOX loaded TriaC-18.

## AUTHOR INFORMATION

## Author Contributions

SM, AB and PS designed the experiments; SM performed synthesis, characterization, rheology study MTT and scratch assay; AB performed the *in vitro* experiments; AS assisted in imaging and flowcytometry analysis; SK and AB^€^ performed the *in-silico* experiments; AG assisted in *in vivo* experiments; PS conceptualized and supervise the project and analyzed the results and wrote the manuscript.

## ACKNOWLEDGMENT

We acknowledge Aftab Hossain Khan for helping us in rheology experiment and data interpretation, Tanvi Agarwal, Int-PhD student of Indian Association for the Cultivation of Science for helping in manuscript writing and editing. Molecular graphics and analyses performed with UCSF Chimera X, developed by the Resource for Biocomputing, Visualization, and Informatics at the University of California, San Francisco, with support from National Institutes of Health R01-GM129325 and the Office of Cyber Infrastructure and Computational Biology, National Institute of Allergy and Infectious Diseases. We acknowledge biorender.com and AI tools for the preparation of schematic representation and for helping in manuscript preparation.

## REFERENCES

(1) Arnold, M.; Singh, D.; Laversanne, M.; Vignat, J.; Vaccarella, S.; Meheus, F.; Cust, A. E.; De Vries, E.; Whiteman, D. C.; Bray, F. Global Burden of Cutaneous Melanoma in 2020 and Projections to 2040. JAMA Dermatology 2022, 158 (5), 495–503.

(2) Zhong, L.; Li, Y.; Xiong, L.; Wang, W.; Wu, M.; Yuan, T.; Yang, W.; Tian, C.; Miao, Z.; Wang, T.; Yang, S. Small Molecules in Targeted Cancer Therapy: Advances, Challenges, and Future Perspectives. Signal Transduction and Targeted Therapy 2021, 6 (1).

(3) Li, J.; Mooney, D. J. Designing Hydrogels for Controlled Drug Delivery. Nat. Rev. Mater. 2016, 1 (12).

(4) Raina, N.; Pahwa, R.; Bhattacharya, J.; Paul, A. K.; Nissapatorn, V.; Pereira, M. de L.; Oliveira, S. M. R.; Dolma, K. G.; Rahmatullah, M.; Wilairatana, P.; Gupta, M. Drug Delivery Strategies and Biomedical Significance of Hydrogels: Translational Considerations. Pharm. 2022, 14 (3), 574.

(5) Rezk, A. I.; Obiweluozor, F. O.; Choukrani, G.; Park, C. H.; Kim, C. S. Drug Release and Kinetic Models of Anticancer Drug (BTZ) from a PH-Responsive Alginate Polydopamine Hydrogel: Towards Cancer Chemotherapy. Int. J. Biol. Macromol. 2019, 141, 388–400.

(6) Takei, T.; Yoshihara, R.; Danjo, S.; Fukuhara, Y.; Evans, C.; Tomimatsu, R.; Ohzuno, Y.; Yoshida, M. Hydrophobically-Modified Gelatin Hydrogel as a Carrier for Charged Hydrophilic Drugs and Hydrophobic Drugs. Int. J. Biol. Macromol. 2020, 149, 140–147.

(7) Ehrbar, M.; Djonov, V. G.; Schnell, C.; Tschanz, S. A.; Martiny-Baron, G.; Schenk, U.; Wood, J.; Burri, P. H.; Hubbell, J. A.; Zisch, A. H. Cell-Demanded Liberation of VEGF121 From Fibrin Implants Induces Local and Controlled Blood Vessel Growth. Circ. Res. 2004, 94 (8), 1124–1132.

(8) Polo Fonseca, L.; Trinca, R. B.; Felisberti, M. I. Amphiphilic Polyurethane Hydrogels as Smart Carriers for Acidic Hydrophobic Drugs. Int. J. Pharm. 2018, 546 (1–2), 106–114.

(9) García-Fernández, L.; Olmeda-Lozano, M.; Benito-Garzón, L.; Pérez-Caballer, A.; San Román, J.; Vázquez-Lasa, B. Injectable Hydrogel-Based Drug Delivery System for Cartilage Regeneration. Mater. Sci. Eng. C 2020, 110, 110702.

(10) Qin, T.; Liao, W.; Yu, L.; Zhu, J.; Wu, M.; Peng, Q.; Han, L.; Zeng, H. Recent Progress in Conductive Self-Healing Hydrogels for Flexible Sensors. J. Polym. Sci. 2022, 60 (18), 2607–2634.

(11) Desai, N. C.; Makwana, A. H.; Senta, R. D. Synthesis, Characterization and Antimicrobial Activity of Some Novel 4-(4-(Arylamino)-6-(Piperidin-1-Yl)-1,3,5-Triazine-2-Ylamino)-N-(Pyrimidin-2-Yl)Benzenesulfonamides. J. Saudi Chem. Soc. 2016, 20 (6), 686–694.

(12) Rambhia, K. J.; Ma, P. X. Controlled Drug Release for Tissue Engineering. J. Control. Release 2015, 219, 119–128.

(13) Keyes, C.; Duhamel, J.; Fung, S. Y.; Bezaire, J.; Chen, P. Self-Assembling Peptide as a Potential Carrier of Hydrophobic Compounds. J. Am. Chem. Soc. 2004, 126 (24), 7522–7532.

(14) Jayawarna, V.; Ali, M.; Jowitt, T. A.; Miller, A. F.; Saiani, A.; Gough, J. E.; Ulijn, R. V. Nanostructured Hydrogels for Three-Dimensional Cell Culture through Self-Assembly of Fluorenylmethoxycarbonyl-Dipeptides. Adv. Mater. 2006, 18 (5), 611–614.

(15) Orbach, R.; Adler-Abramovich, L.; Zigerson, S.; Mironi-Harpaz, I.; Seliktar, D.; Gazit, E. Self-Assembled Fmoc-Peptides as a Platform for the Formation of Nanostructures and Hydrogels. Biomacromolecules 2009, 10 (9), 2646–2651.

(16) Zhang, Y.; Gu, H.; Yang, Z.; Xu, B. Supramolecular Hydrogels Respond to Ligand-Receptor Interaction. J. Am. Chem. Soc. 2003, 125 (45), 13680–13681.

(17) Hartgerink, J. D.; Beniash, E.; Stupp, S. I. Self-Assembly and Mineralization of Peptide-Amphiphile Nanofibers. Science (80-. ). 2001, 294 (5547), 1684–1688.

(18) Sanborn, T. J.; Messersmith, P. B.; Barron, A. E. In Situ Crosslinking of a Biomimetic Peptide-PEG Hydrogel via Thermally Triggered Activation of Factor XIII. Biomaterials 2002, 23, 2703–2710.

(19) Ma, X.; Xu, T.; Chen, W.; Qin, H.; Chi, B.; Ye, Z. Injectable Hydrogels Based on the Hyaluronic Acid and Poly (γ-Glutamic Acid) for Controlled Protein Delivery. Carbohydr. Polym. 2018, 179, 100–109.

(20) Caballero Aguilar, L. M.; Silva, S. M.; Moulton, S. E. Growth Factor Delivery: Defining the next Generation Platforms for Tissue Engineering. J. Control. Release 2019, 306, 40–58.

(21) Wang, P.; Huang, S.; Hu, Z.; Yang, W.; Lan, Y.; Zhu, J.; Hancharou, A.; Guo, R.; Tang, B. In Situ Formed Anti-Inflammatory Hydrogel Loading Plasmid DNA Encoding VEGF for Burn Wound Healing. Acta Biomater. 2019, 100, 191–201.

(22) Chalanqui, M. J.; Pentlavalli, S.; McCrudden, C.; Chambers, P.; Ziminska, M.; Dunne, N.; McCarthy, H. O. Influence of Alginate Backbone on Efficacy of Thermo-Responsive Alginate-g-P(NIPAAm) Hydrogel as a Vehicle for Sustained and Controlled Gene Delivery. Mater. Sci. Eng. C 2019, 95, 409–421.

(23) Tao, G.; Wang, Y.; Cai, R.; Chang, H.; Song, K.; Zuo, H.; Zhao, P.; Xia, Q.; He, H. Design and Performance of Sericin/Poly(Vinyl Alcohol) Hydrogel as a Drug Delivery Carrier for Potential Wound Dressing Application. Mater. Sci. Eng. C 2019, 101, 341–351.

(24) Hu, J.; Zhang, Y.; Tang, N.; Lu, Y.; Guo, P.; Huang, Z. Discovery of Novel 1,3,5-Triazine Derivatives as Potent Inhibitor of Cervical Cancer via Dual Inhibition of PI3K/MTOR. Bioorg. Med. Chem. 2021, 32, 115997.

(25) Poulsen, A.; Williams, M.; Nagaraj, H. M.; William, A. D.; Wang, H.; Soh, C. K.; Xiong, Z. C.; Dymock, B. Structure-Based Optimization of Morpholino-Triazines as PI3K and MTOR Inhibitors. Bioorg. Med. Chem. Lett. 2012, 22 (2), 1009–1013.

(26) Sakakibara, N.; Balboni, G.; Congiu, C.; Onnis, V.; Demizu, Y.; Misawa, T.; Kurihara, M.; Kato, Y.; Maruyama, T.; Toyama, M.; Okamoto, M.; Baba, M. Design, Synthesis, and Anti-HIV-1 Activity of 1-Substituted 3-(3,5-Dimethylbenzyl)Triazine Derivatives. Antivir. Chem. Chemother. 2015, 24 (2), 62–71.

(27) Chen, X.; Meng, Q.; Qiu, L.; Zhan, P.; Liu, H.; De Clercq, E.; Pannecouque, C.; Liu, X. Design, Synthesis, and Anti-HIV Evaluation of Novel Triazine Derivatives Targeting the Entrance Channel of the NNRTI Binding Pocket. Chem. Biol. Drug Des. 2015, 86 (1), 122–128.

(28) Zhang, Q.; Peng, Y.; Wang, X. I.; Keenan, S. M.; Arora, S.; Welsh, W. J. Highly Potent Triazole-Based Tubulin Polymerization Inhibitors. J. Med. Chem. 2007, 50 (4), 749–754.

(29) Pogorelčnik, B.; Janežič, M.; Sosič, I.; Gobec, S.; Solmajer, T.; Perdih, A. 4,6-Substituted-1,3,5-Triazin-2(1H)-Ones as Monocyclic Catalytic Inhibitors of Human DNA Topoisomerase IIα Targeting the ATP Binding Site. Bioorg. Med. Chem. 2015, 23 (15), 4218–4229.

(30) Zhang, C.; Zhang, J.; Qin, Y.; Song, H.; Huang, P.; Wang, W.; Wang, C.; Li, C.; Wang, Y.; Kong, D. Co-Delivery of Doxorubicin and Pheophorbide A by Pluronic F127 Micelles for Chemo-Photodynamic Combination Therapy of Melanoma. J. Mater. Chem. B 2018, 6 (20), 3305–3314.

(31) Rageot, D.; Bohnacker, T.; Keles, E.; McPhail, J. A.; Hoffmann, R. M.; Melone, A.; Borsari, C.; Sriramaratnam, R.; Sele, A. M.; Beaufils, F.; Hebeisen, P.; Fabbro, D.; Hillmann, P.; Burke, J. E.; Wymann, M. P. (S)-4-(Difluoromethyl)-5-(4-(3-Methylmorpholino)-6-Morpholino-1,3,5-Triazin-2-Yl)Pyridin-2-Amine (PQR530), a Potent, Orally Bioavailable, and Brain-Penetrable Dual Inhibitor of Class i PI3K and MTOR Kinase. J. Med. Chem. 2019, 62 (13), 6241–6261.

(32) Berman, H. M.; Westbrook, J.; Feng, Z.; Gilliland, G.; Bhat, T. N.; Weissig, H.; Shindyalov, I. N.; Bourne, P. E. The Protein Data Bank. Nucleic Acids Res. 2000, 28 (1), 235–242.

(33) Šali, A.; Blundell, T. L. Comparative Protein Modelling by Satisfaction of Spatial Restraints. J. Mol. Biol. 1993, 234 (3), 779–815.

(34) Colovos, C.; Yeates, T. O. Verification of Protein Structures: Patterns of Nonbonded Atomic Interactions. Protein Sci. 1993, 2 (9), 1511–1519.

(35) Ramachandran, S.; Kota, P.; Ding, F.; Dokholyan, N. V. Automated Minimization of Steric Clashes in Protein Structures. Proteins Struct. Funct. Bioinforma. 2011, 79 (1), 261–270.

(36) Sanner, M. Python: A Programming Language for Software Integration and Development. J. Mol. Graph. Model. 1999.

(37) Morris, G. M.; Ruth, H.; Lindstrom, W.; Sanner, M. F.; Belew, R. K.; Goodsell, D. S.; Olson, A. J. AutoDock4 and AutoDockTools4: Automated Docking with Selective Receptor Flexibility. J. Comput. Chem. 2009, 30 (16), 2785.

(38) Meng, E. C.; Goddard, T. D.; Pettersen, E. F.; Couch, G. S.; Pearson, Z. J.; Morris, J. H.; Ferrin, T. E. UCSF ChimeraX: Tools for Structure Building and Analysis. Protein Sci. 2023, 32 (11), e4792.

(39) Adasme, M. F.; Linnemann, K. L.; Bolz, S. N.; Kaiser, F.; Salentin, S.; Haupt, V. J.; Schroeder, M. PLIP 2021: Expanding the Scope of the Protein–Ligand Interaction Profiler to DNA and RNA. Nucleic Acids Res. 2021, 49 (W1), W530–W534.

(40) Jorgensen, W. L.; Chandrasekhar, J.; Madura, J. D.; Impey, R. W.; Klein, M. L. Comparison of Simple Potential Functions for Simulating Liquid Water. J. Chem. Phys. 1983, 79 (2), 926–935.

(41) Vanommeslaeghe, K.; Hatcher, E.; Acharya, C.; Kundu, S.; Zhong, S.; Shim, J.; Darian, E.; Guvench, O.; Lopes, P.; Vorobyov, I.; Mackerell, A. D. CHARMM General Force Field: A Force Field for Drug-like Molecules Compatible with the CHARMM All-Atom Additive Biological Force Fields. J. Comput. Chem. 2010, 31 (4), 671–690.

(42) MacKerell, A. D.; Bashford, D.; Bellott, M.; Dunbrack, R. L.; Evanseck, J. D.; Field, M. J.; Fischer, S.; Gao, J.; Guo, H.; Ha, S.; Joseph-McCarthy, D.; Kuchnir, L.; Kuczera, K.; Lau, F. T. K.; Mattos, C.; Michnick, S.; Ngo, T.; Nguyen, D. T.; Prodhom, B.; Reiher, W. E.; Roux, B.; Schlenkrich, M.; Smith, J. C.; Stote, R.; Straub, J.; Watanabe, M.; Wiórkiewicz-Kuczera, J.; Yin, D.; Karplus, M. All-Atom Empirical Potential for Molecular Modeling and Dynamics Studies of Proteins. J. Phys. Chem. B 1998, 102 (18), 3586–3616.

(43) Brooks, B. R.; Bruccoleri, R. E.; Olafson, B. D.; States, D. J.; Swaminathan, S.; Karplus, M. CHARMM: A Program for Macromolecular Energy, Minimization, and Dynamics Calculations. J. Comput. Chem. 1983, 4 (2), 187–217.

(44) Phillips, J. C.; Braun, R.; Wang, W.; Gumbart, J.; Tajkhorshid, E.; Villa, E.; Chipot, C.; Skeel, R. D.; Kalé, L.; Schulten, K. Scalable Molecular Dynamics with NAMD. J. Comput. Chem. 2005, 26 (16), 1781–1802.

(45) Darden, T.; York, D.; Pedersen, L. Particle Mesh Ewald: An NLog(N) Method for Ewald Sums in Large Systems. J. Chem. Phys. 1993, 98, 5648.

(46) Feller, S. E.; Zhang, Y.; Pastor, R. W.; Brooks, B. R. Constant Pressure Molecular Dynamics Simulation: The Langevin Piston Method. J. Chem. Phys. 1995, 103 (11), 4613–4621.

(47) Martyna, G. J.; Tobias, D. J.; Klein, M. L. Constant Pressure Molecular Dynamics Algorithms. J. Chem. Phys. 1994, 101 (5), 4177–4189.

(48) Evans, D. J.; Holian, B. L. The Nose–Hoover Thermostat. J. Chem. Phys. 1985, 83 (8), 4069–4074.

(49) Martyna, G. J.; Klein, M. L.; Tuckerman, M. Nosé–Hoover Chains: The Canonical Ensemble via Continuous Dynamics. J. Chem. Phys. 1992, 97 (4), 2635–2643.

(50) Hung, P. K.; Vinh, L. T.; Nghiep, D. M.; Nguyen, P. N. Computer Simulation of Liquid Al2O3. Journal of Physics Condensed Matter. 2006, 18 (41), 9309–9322.

(51) Jiang, H.; Duan, L.; Ren, X.; Gao, G. Hydrophobic Association Hydrogels with Excellent Mechanical and Self-Healing Properties. Eur. Polym. J. 2019, 112, 660–669.

(52) Zhu, L.; Lu, Q.; Bian, T.; Yang, P.; Yang, Y.; Zhang, L. Fabrication and Characterization of π-π Stacking Peptide-Contained Double Network Hydrogels. ACS Biomater. Sci. Eng. 2023, 9 (8), 4761–4769.

(53) Sarkar, K.; Dastidar, P. Supramolecular Hydrogel Derived from a C3-Symmetric Boronic Acid Derivative for Stimuli-Responsive Release of Insulin and Doxorubicin. Langmuir 2018, 34 (2), 685–692.

(54) Del Campo, J. M.; Birrer, M.; Davis, C.; Fujiwara, K.; Gollerkeri, A.; Gore, M.; Houk, B.; Lau, S.; Poveda, A.; González-Martín, A.; Muller, C.; Muro, K.; Pierce, K.; Suzuki, M.; Vermette, J.; Oza, A. A Randomized Phase II Non-Comparative Study of PF-04691502 and Gedatolisib (PF-05212384) in Patients with Recurrent Endometrial Cancer. Gynecol. Oncol. 2016, 142 (1), 62–69.

(55) Martín, M.; Chan, A.; Dirix, L.; O’Shaughnessy, J.; Hegg, R.; Manikhas, A.; Shtivelband, M.; Krivorotko, P.; Batista López, N.; Campone, M.; Ruiz Borrego, M.; Khan, Q. J.; Beck, J. T.; Ramos Vázquez, M.; Urban, P.; Goteti, S.; Di Tomaso, E.; Massacesi, C.; Delaloge, S. A Randomized Adaptive Phase II/III Study of Buparlisib, a Pan-Class I PI3K Inhibitor, Combined with Paclitaxel for the Treatment of HER2– Advanced Breast Cancer (BELLE-4). Ann. Oncol. 2017, 28 (2), 313–320.

(56) Massacesi, C.; di Tomaso, E.; Urban, P.; Germa, C.; Quadt, C.; Trandafir, L.; Aimone, P.; Fretault, N.; Dharan, B.; Tavorath, R.; Hirawat, S. PI3K Inhibitors as New Cancer Therapeutics: Implications for Clinical Trial Design. Onco. Targets. Ther. 2016, 9, 203–210.

(57) Venkatesan, A. M.; Dehnhardt, C. M.; Delos Santos, E. D.; Chen, Z.; Dos Santos, O. D.; Ayral-Kaloustian, S.; Khafizova, G.; Brooijmans, N.; Mallon, R.; Hollander, I.; Feldberg, L.; Lucas, J.; Yu, K.; Gibbons, J.; Abraham, R. T.; Chaudhary, I.; Mansour, T. S. Bis(Morpholino-l,3,5-Triazine) Derivatives: Potent Adenosine 5′-Triphosphate Competitive Phosphatidylinositol-3-Kinase/Mammalian Target of Rapamycin Inhibitors: Discovery of Compound 26 (PKI-587), a Highly Efficacious Dual Inhibitor. J. Med. Chem. 2010, 53 (6), 2636–2645.

(58) Zhu, J.; Li, K.; Yu, L.; Chen, Y.; Cai, Y.; Jin, J.; Hou, T. Targeting Phosphatidylinositol 3-Kinase Gamma (PI3Kγ): Discovery and Development of Its Selective Inhibitors. 2020. 41 (3), 1599–1621.

(59) Zhu, J.; Li, K.; Yu, L.; Chen, Y.; Cai, Y.; Jin, J.; Hou, T. Targeting Phosphatidylinositol 3-Kinase Gamma (PI3Kγ): Discovery and Development of Its Selective Inhibitors. 2020. 41 (3), 1599–1621.

(60) Venkatesan, A. M.; Dehnhardt, C. M.; Delos Santos, E. D.; Chen, Z.; Dos Santos, O. D.; Ayral-Kaloustian, S.; Khafizova, G.; Brooijmans, N.; Mallon, R.; Hollander, I.; Feldberg, L.; Lucas, J.; Yu, K.; Gibbons, J.; Abraham, R. T.; Chaudhary, I.; Mansour, T. S. Bis(Morpholino-l,3,5-Triazine) Derivatives: Potent Adenosine 5′-Triphosphate Competitive Phosphatidylinositol-3-Kinase/Mammalian Target of Rapamycin Inhibitors: Discovery of Compound 26 (PKI-587), a Highly Efficacious Dual Inhibitor. J. Med. Chem. 2010, 53 (6), 2636–2645.

(61) Konar, S.; Sinha, S. K.; Datta, S.; Ghorai, P. K. Probing the Dynamics between the Substrate and the Product towards Glucose Tolerance of Halothermothrix Orenii β-Glucosidase. J. Biomol. Struct. Dyn. 2021, 39 (15), 5438–5448.

