## Supplementary information for "Design and Synthesis of Triazine-Based Hydrogel for Combined Targeted Doxorubicin Delivery and PI3K Inhibition"

$ Equal Contribution

#Equal Contribution

Table of Contents

1.1^1^H NMR Spectra of **TriaC-12**	**S1**

1.2.^13^C NMR Spectra of **TriaC-12**	**S2**

1.3. MALDI-TOF Spectra of **TriaC-12**	**S3**

2.1. ^1^H NMR Spectra of **TriaC-14**	**S4**

2.2. ^13^C NMR Spectra of **TriaC-14**	**S5**

2.3. MALDI-TOF Spectra of **TriaC-14**	**S6**

3.1. ^1^H NMR Spectra of **TriaC-16**	**S7**

3.2. ^13^C NMR Spectra of **TriaC-16**	**S8**

3.3. MALDI-TOF Spectra of **TriaC-16**	**S9**

4.1. ^1^H NMR Spectra of **TriaC-18**	**S10**

4.2.^13^C NMR Spectra of **TriaC-18**	**S11**

4.3. MALDI-TOF Spectra of **TriaC-18**	**S12**

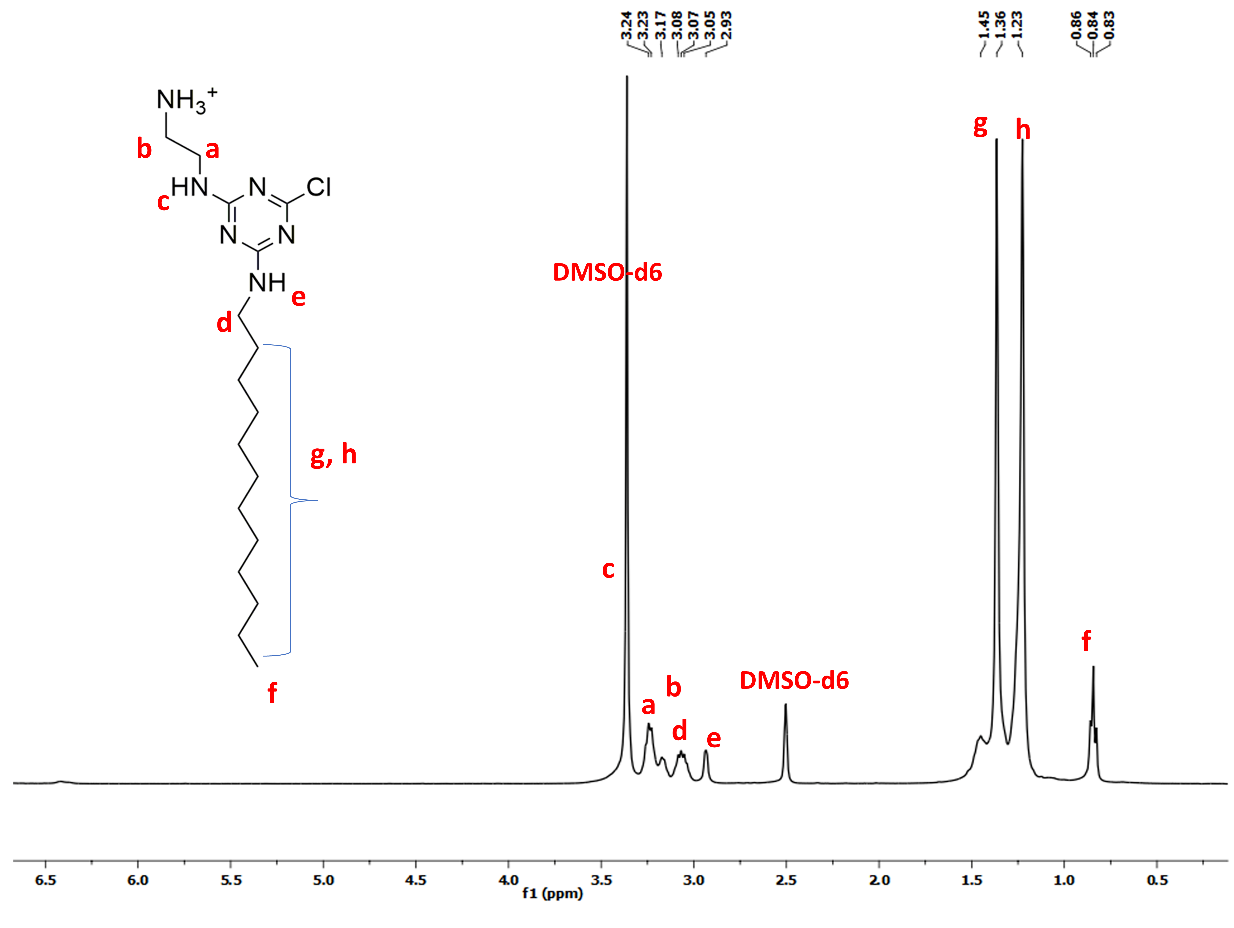

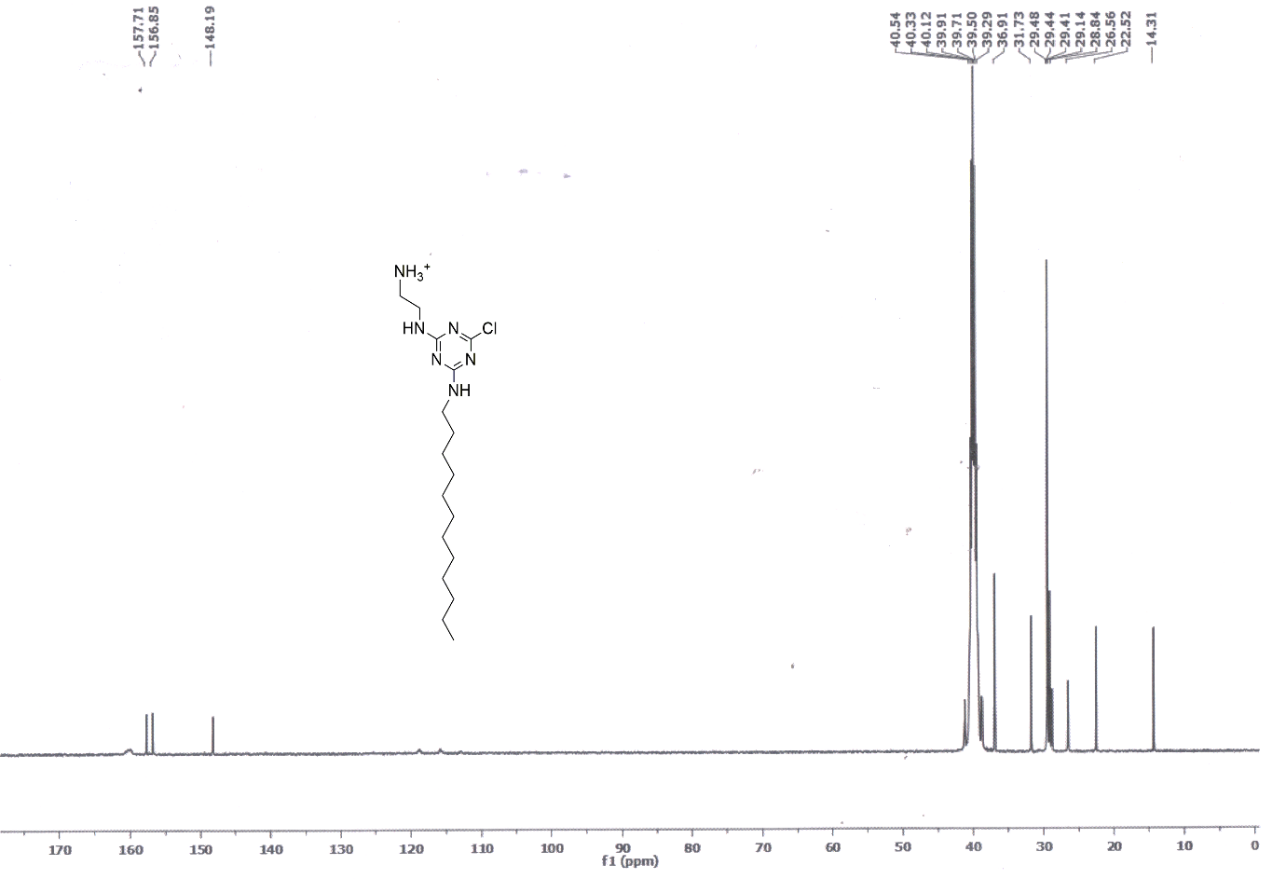
**Figure S1:** ^1^H NMR of **TriaC-12**

**Figure S2:** ^13^C NMR of **TriaC-12**

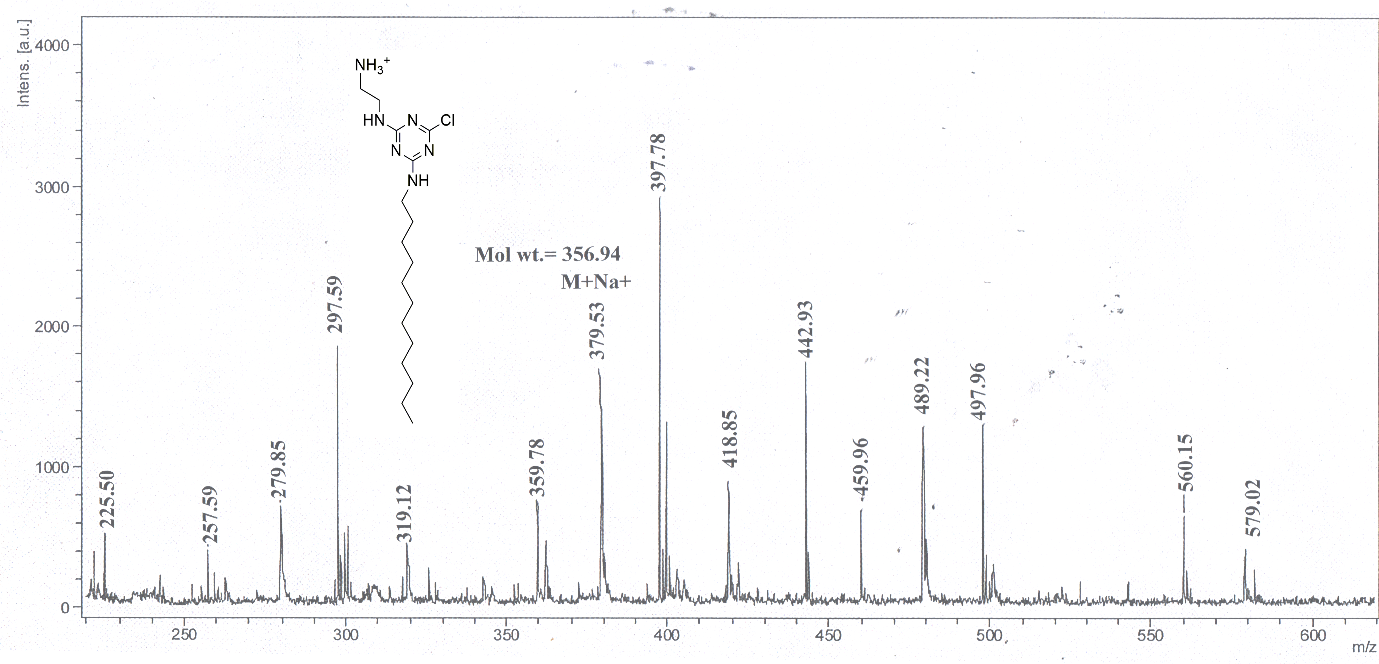

**Figure S3:** MALDI-TOF Spectra of **TriaC-12**

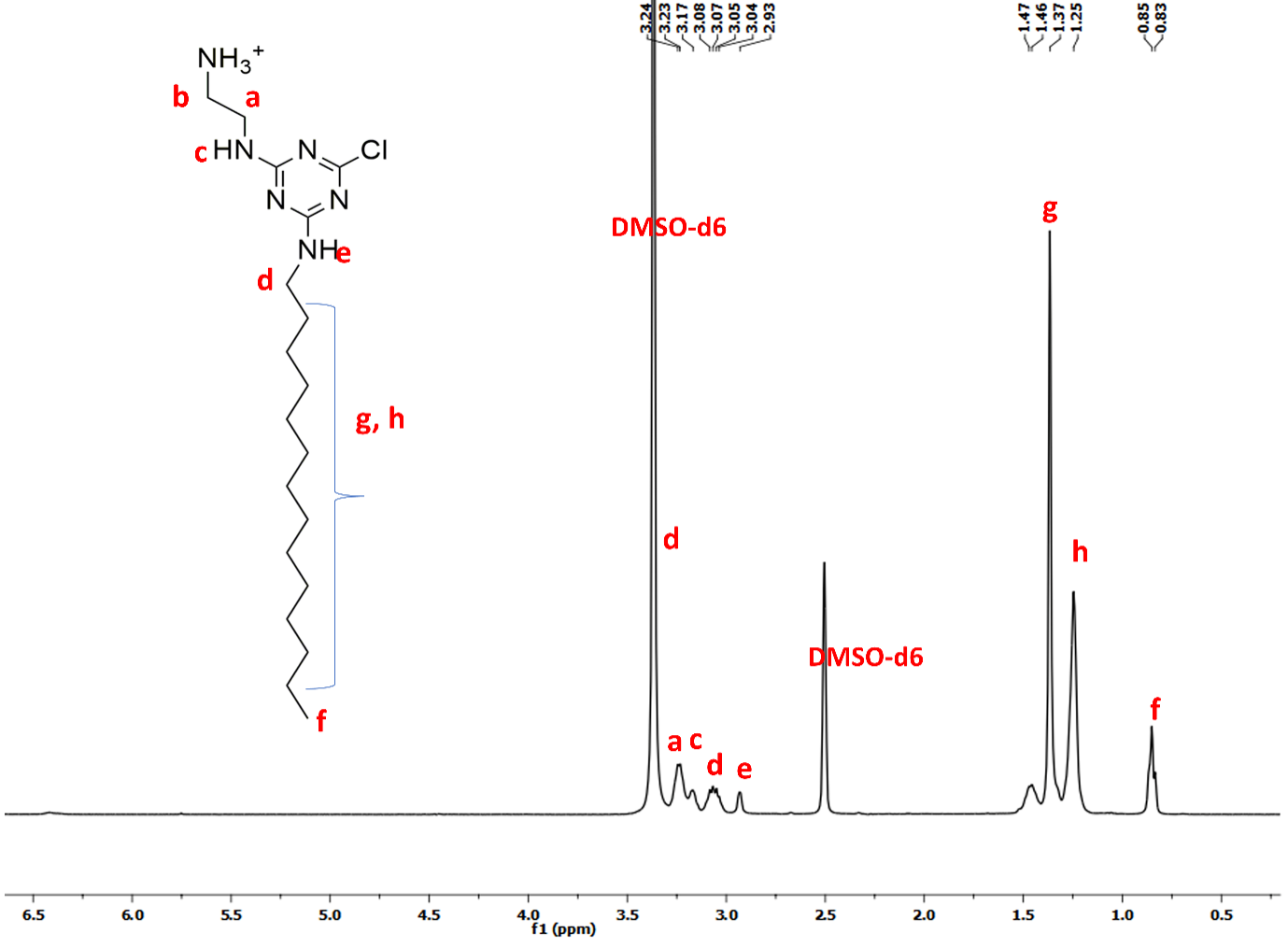

**Figure S4:** ^1^H NMR of **TriaC-14**

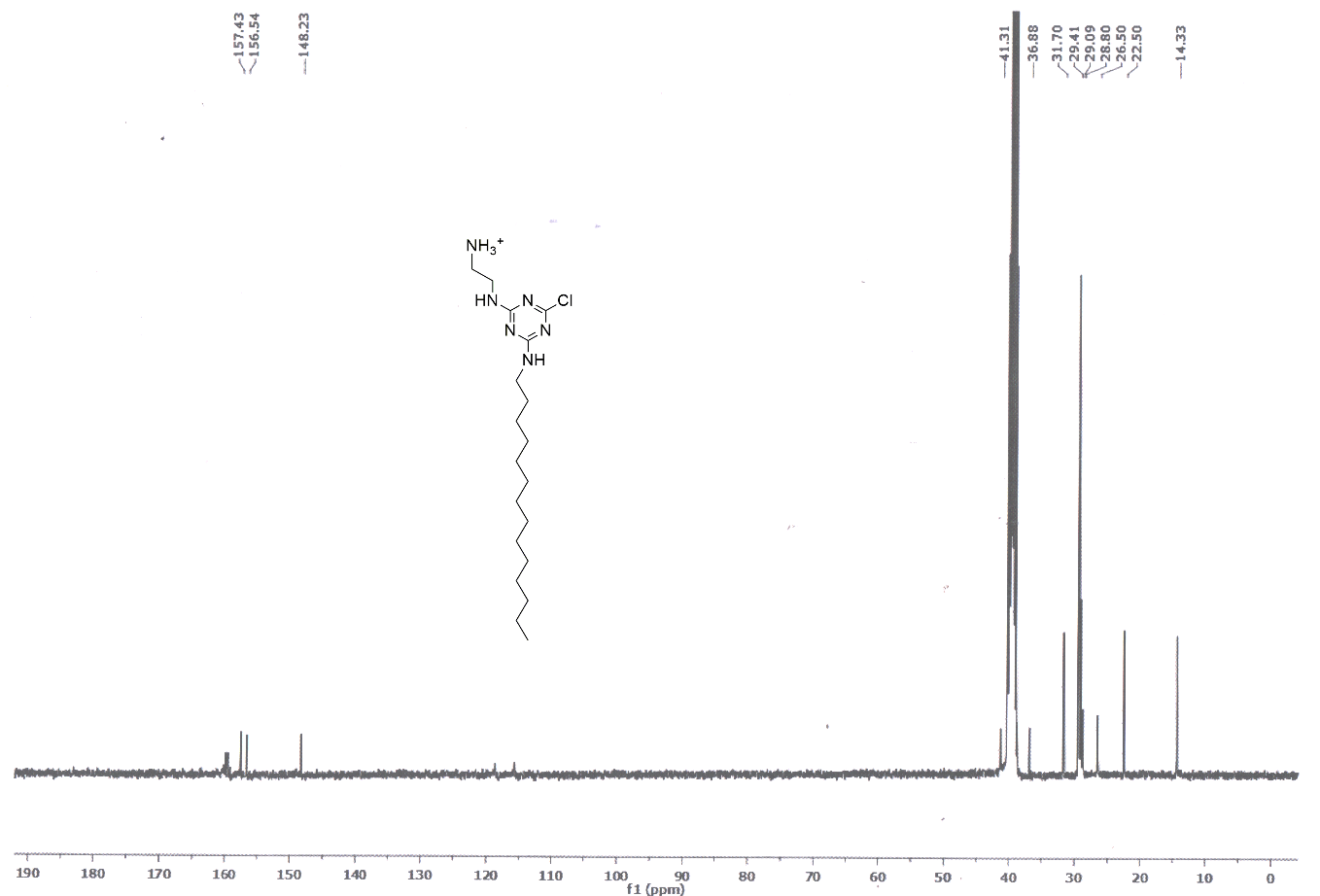

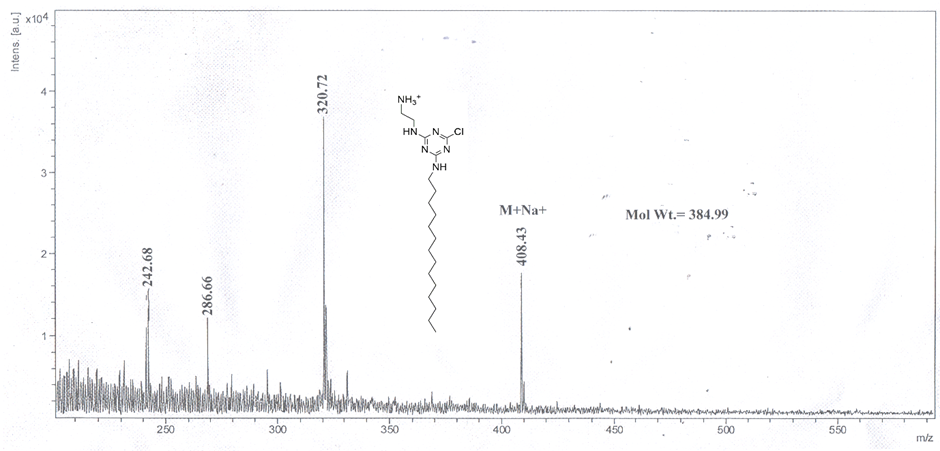
**Figure S5:** ^13^C NMR of **TriaC-14**

**Figure S6:** MALDI-TOF Spectra of **TriaC-14**

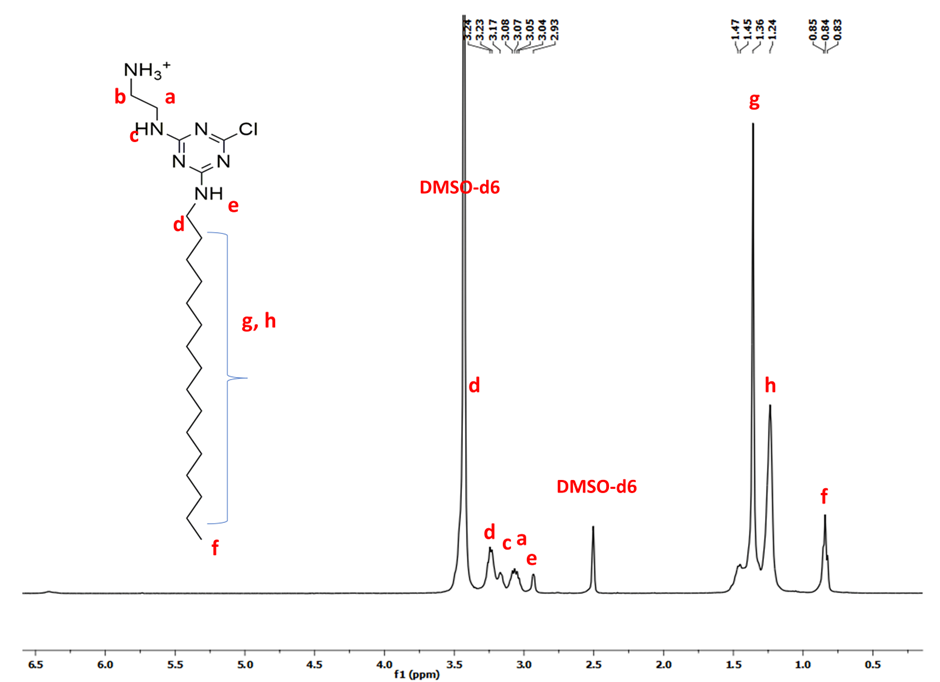

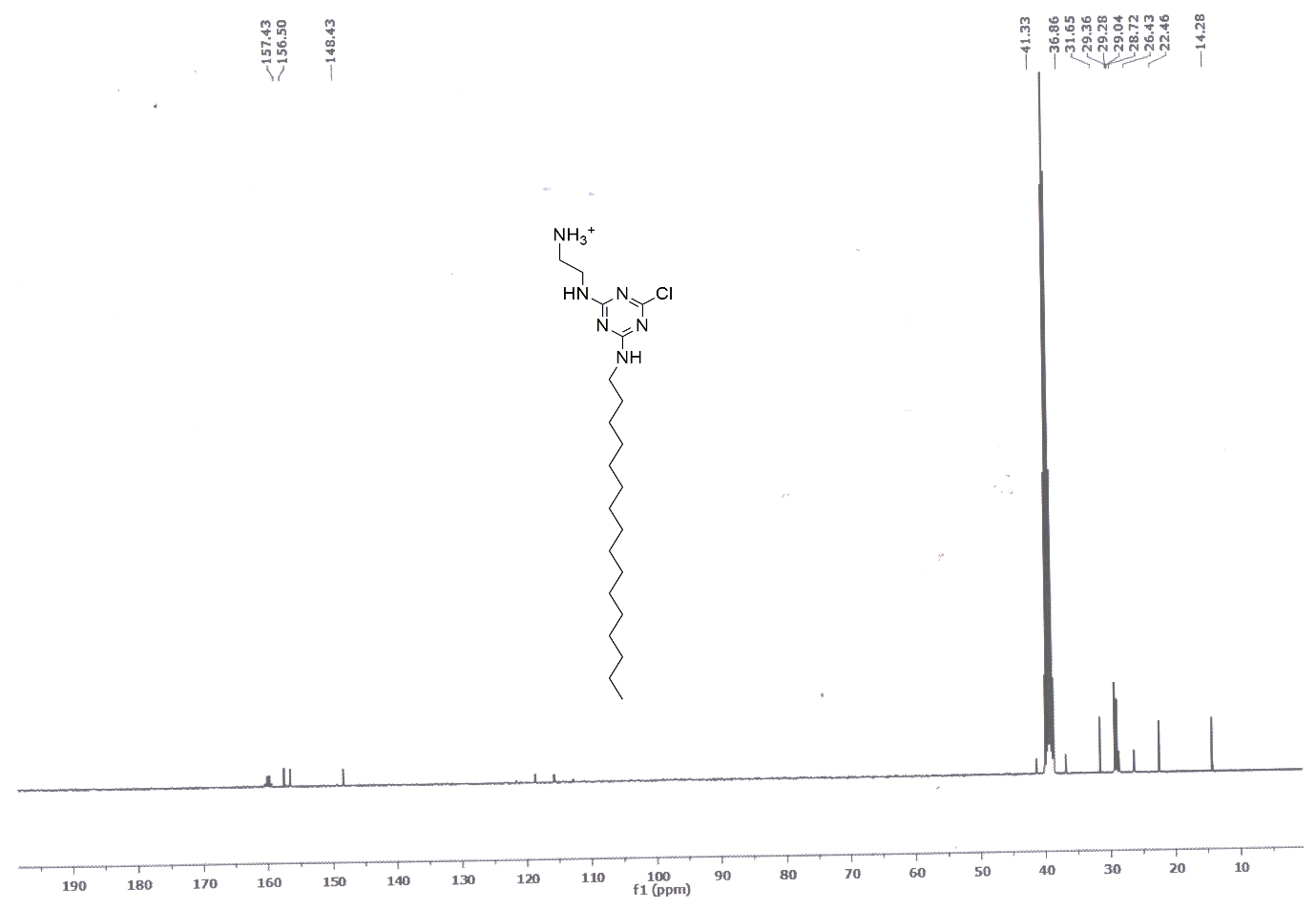
**Figure S7:** ^1^H NMR of **TriaC-16**

**Figure S8:** ^13^C NMR of **TriaC-16**

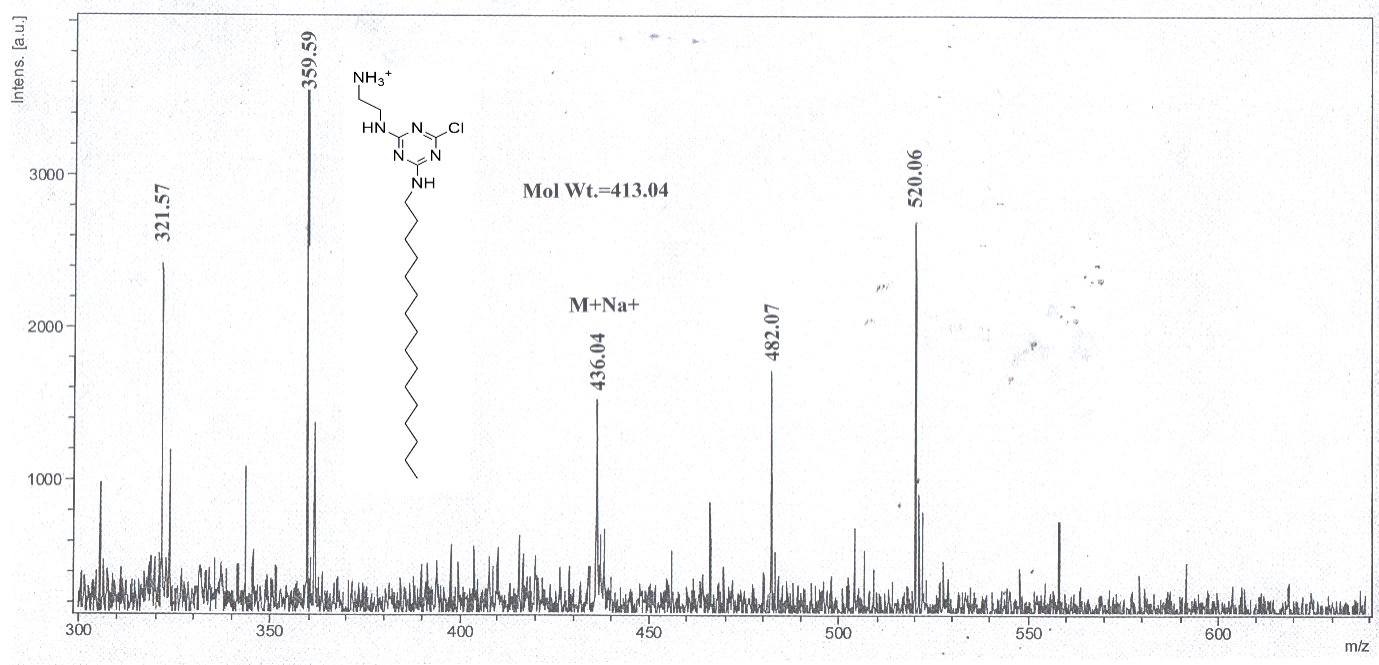

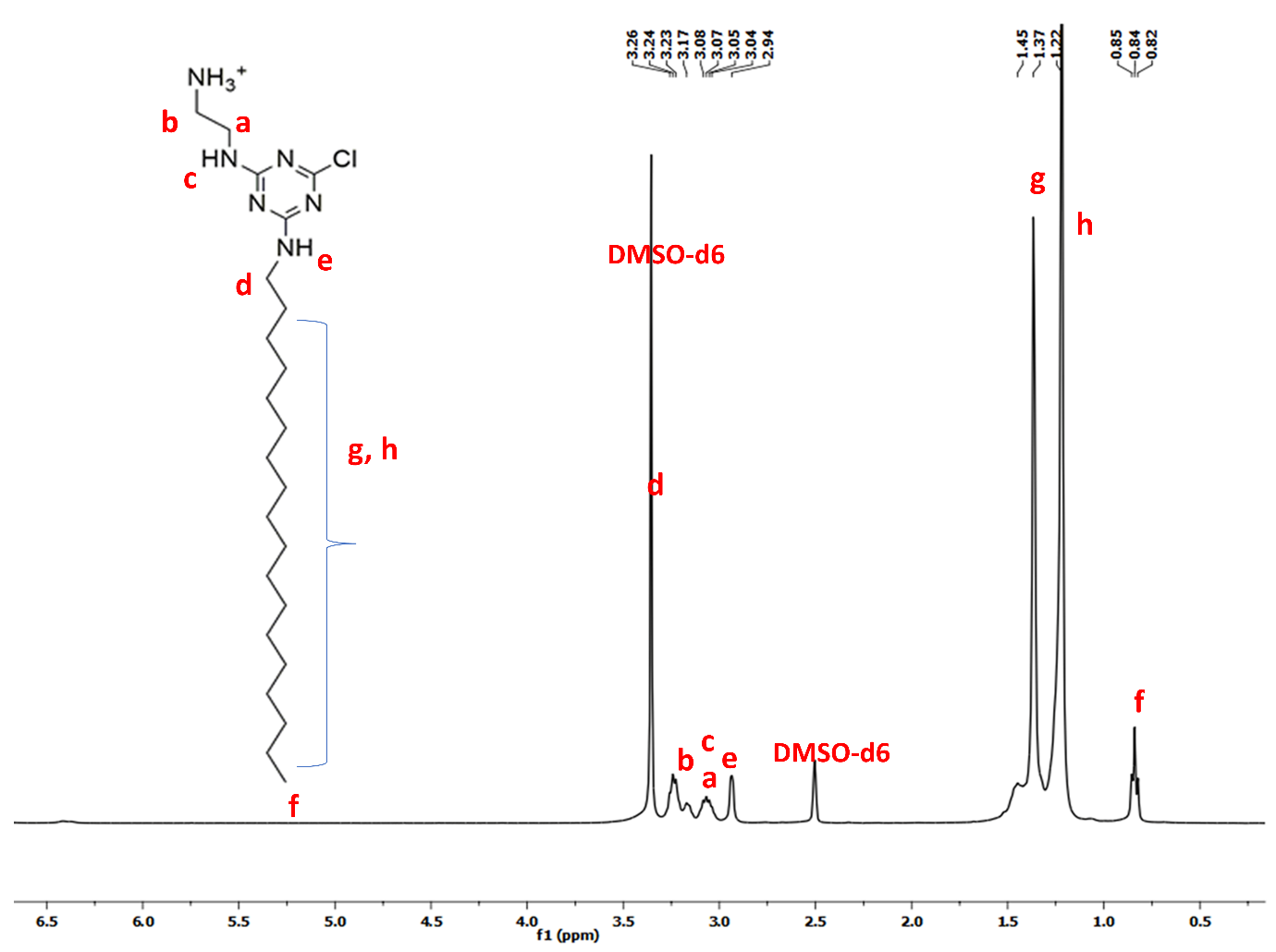
**Figure S9:** MALDI-TOF Spectra of **TriaC-16**

**Figure S10:** ^1^H NMR of **TriaC-18**

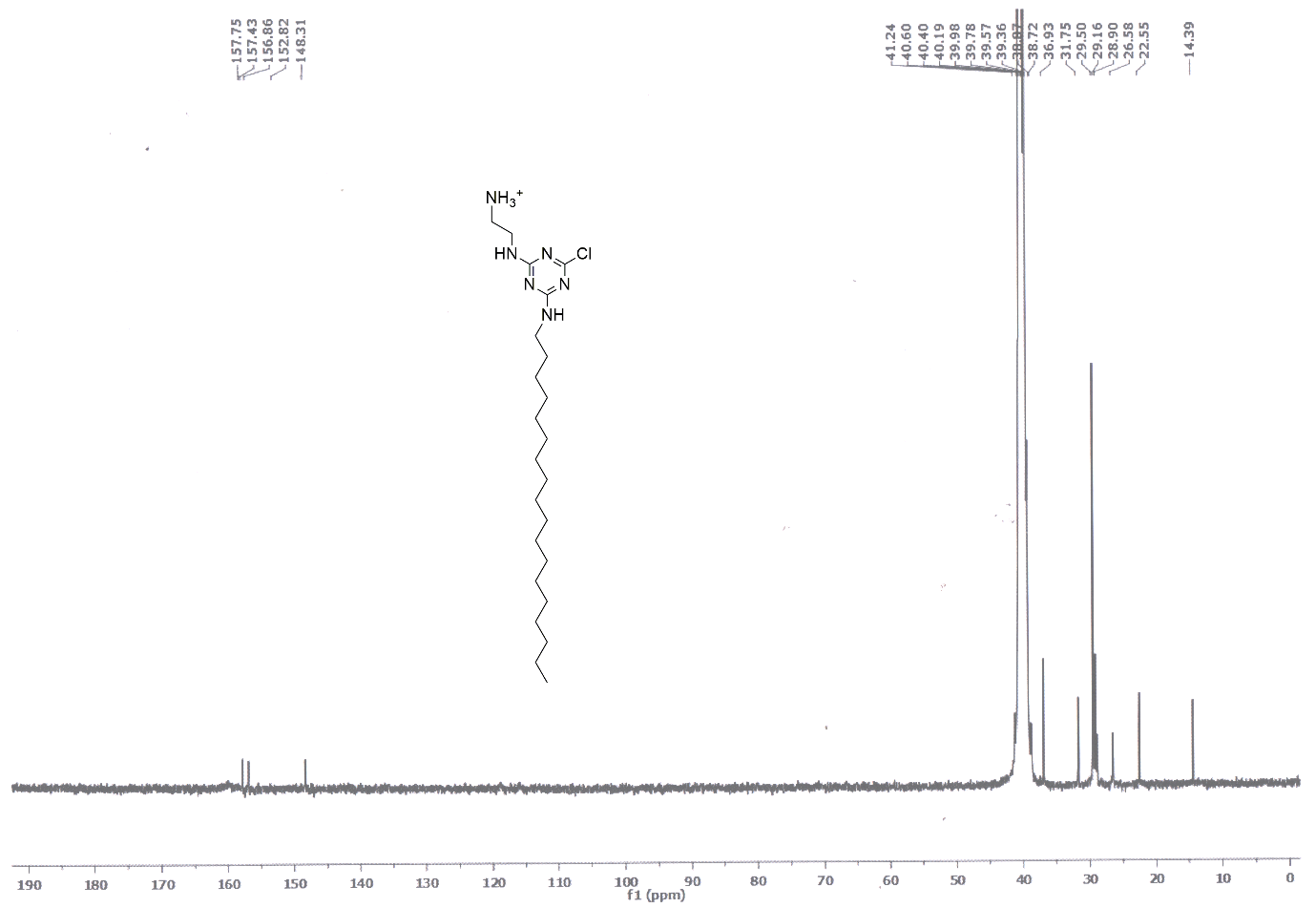

**Figure S11:** ^13^C NMR of **TriaC-18**

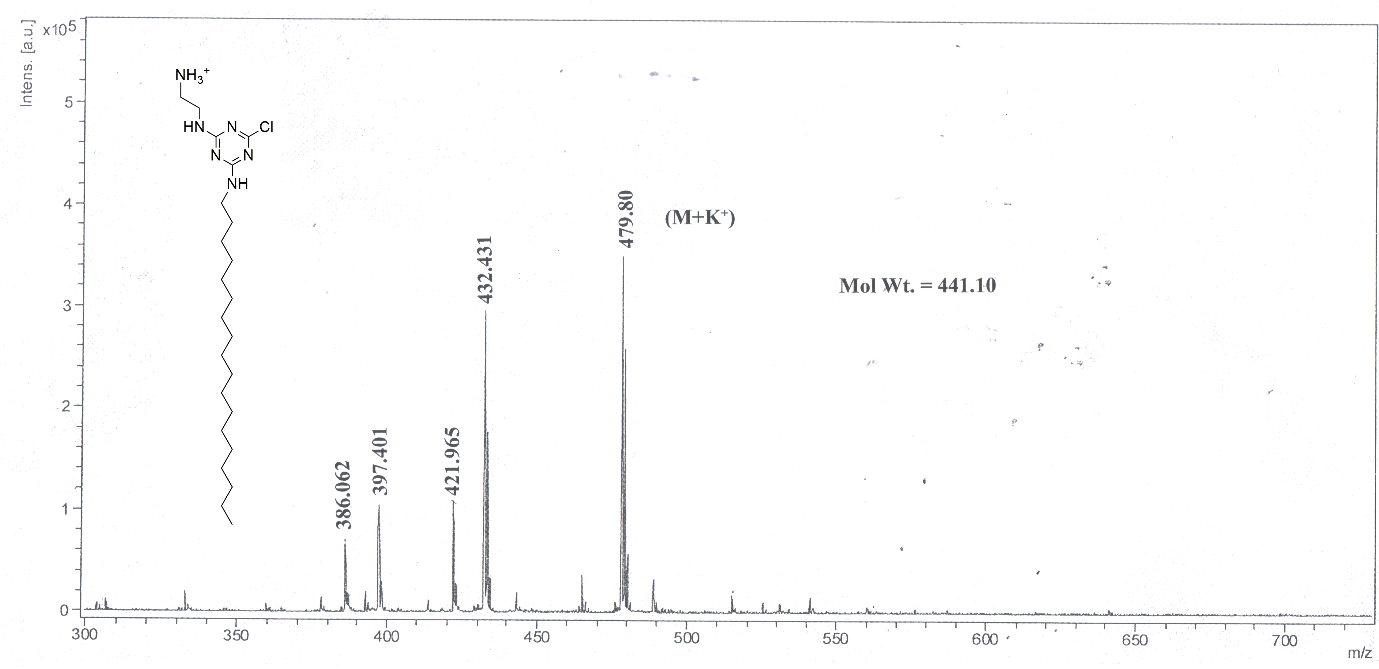

**Figure S12:** MALDI-TOF Spectra of **TriaC-18**

**Figure S13: (A)**Averag
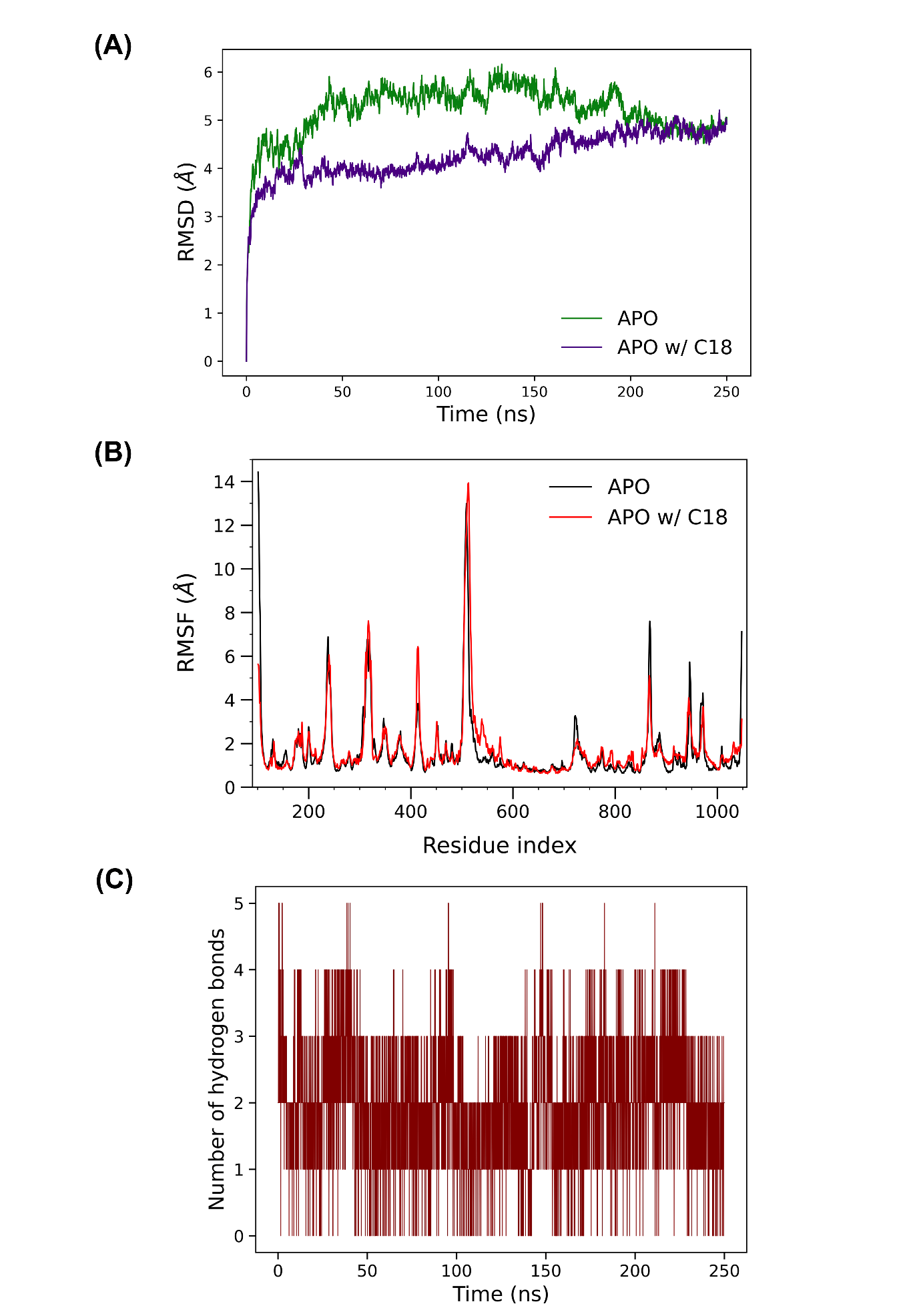
eRMSD profile of two replicates of control (APO) and **TriaC-18**bound PI3K catalytic subunit p110α for 250 ns **(B)** RMSF profile of the **TriaC-18** bound complex (APO w/ C18), along with the control (APO), whole protein **(C)**Hydrogen bond distribution profile showing the hydrogen bond count per frame between the protein and ligand for the entire simulation period of 250 ns.

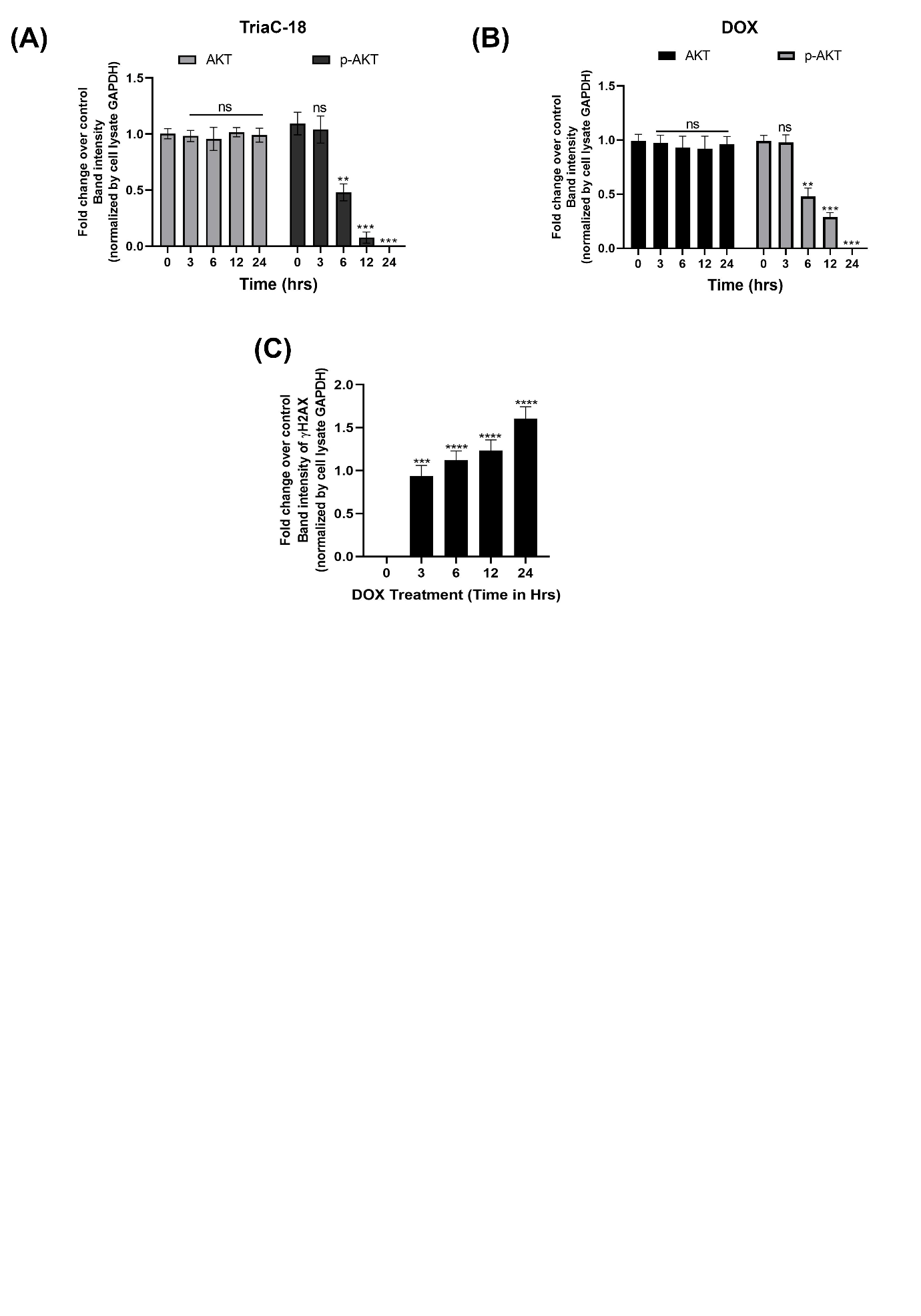
**Figure S14: (A, B)**Quantification graph of western blot analysis of AKT phosphorylation along with total AKT expression in different time point (0-24 hrs.) after treating A375 cells with**TriaC-18**. **(C)**Quantification graph of western blot analysis ofdsDNA break marker γ-H2AX expression in A375 cells in different time point (0-24 hrs) after doxorubicintreatment.

**Table 1: List of Antibodies**

| Antibodies | Final Dilution used | Company and catalog details |
| --- | --- | --- |
| AKT | 1:1000 | Biorad, AbD30769 |
| p-AKT | 1:1000 | CST#9271 |
| FOXO3a | 1:1000 | A0102 |
| p-FOXO3a | 1:1000 | CST#9466 |
| Bim | 1:1000 | A0295 |
| PUMA | 1:1000 | A17138 |
| NOXA | 1:1000 | PA5-19977 |
| Bax | 1:1000 | PA5-85918 |
| Bcl2 | 1:1000 | PA5-27094 |
| Cleaved PARP | 1:1000 | CST#9541 |
| Cleaved Caspase3 | 1:1000 | MAB835 |
| γH2AX | 1:1000 | A11412 |
| HRP-tagged secondary ab (mouse) | 1:10000 | Sigma (A9917) |
| HRP-tagged secondary ab (Rabbit) | 1:10000 | Sigma (A6154) |

**Table 2: Average RMSD of the two systems: control (APO) and the TriaC-18-boundprotein
(APO w/ C18)**

|  | APO | APO w/ C18 |
| --- | --- | --- |
| Average RMSD for last 100 ns | 4.676 Å | 5.132 Å |

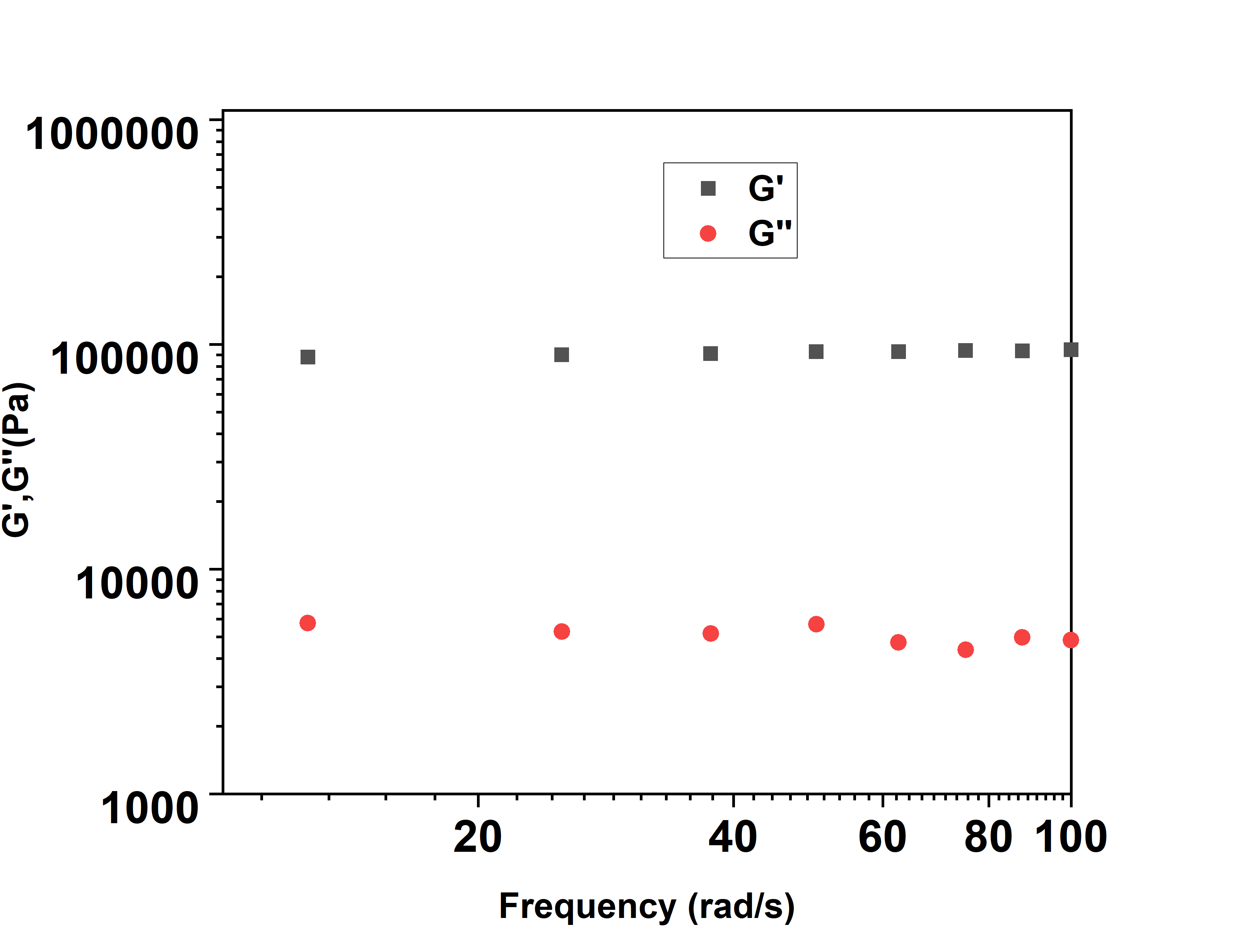

**Figure S15.** tan delta (G’’/G’)

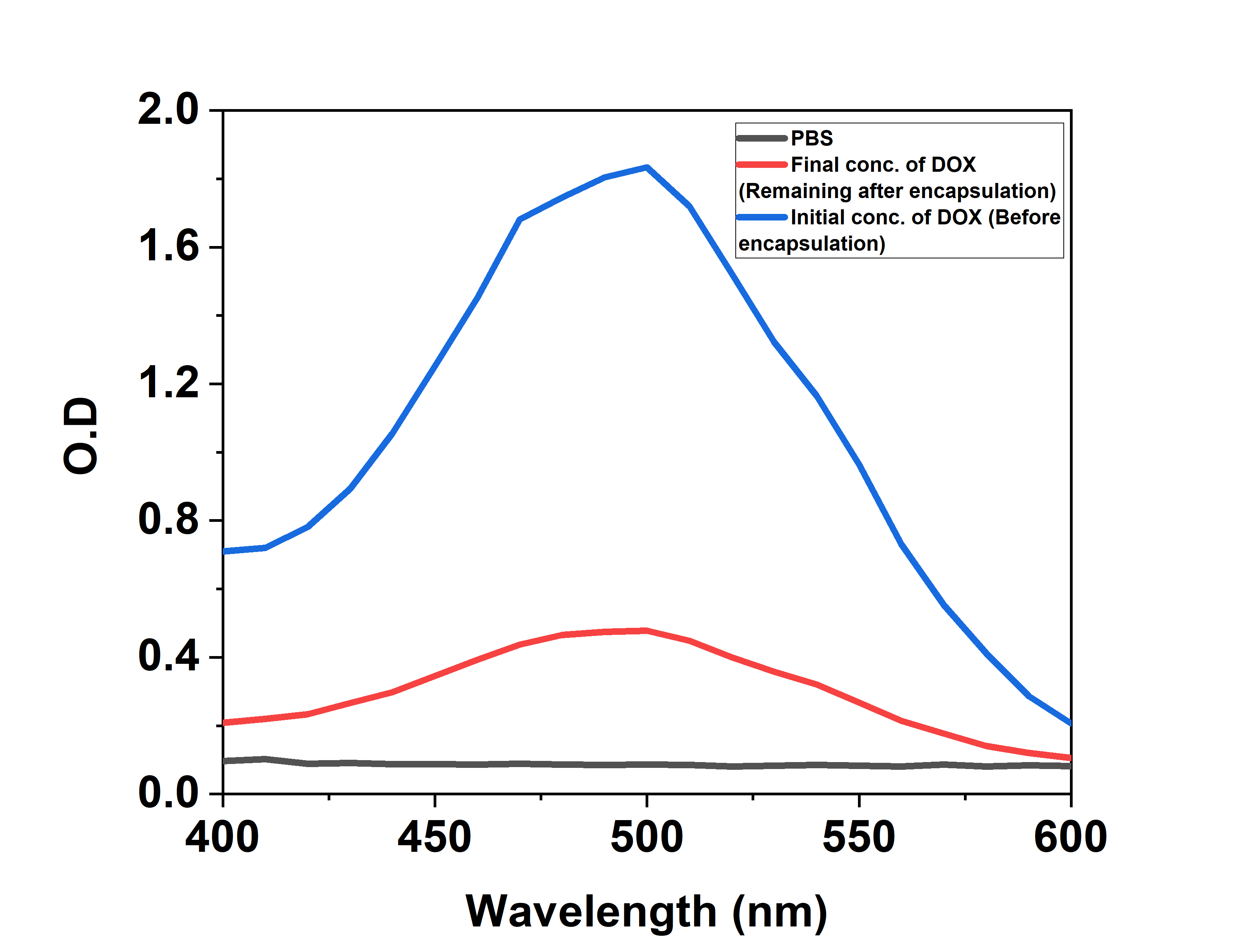

**Figure S16.** DOX Loading capacity of **TriaC-18**

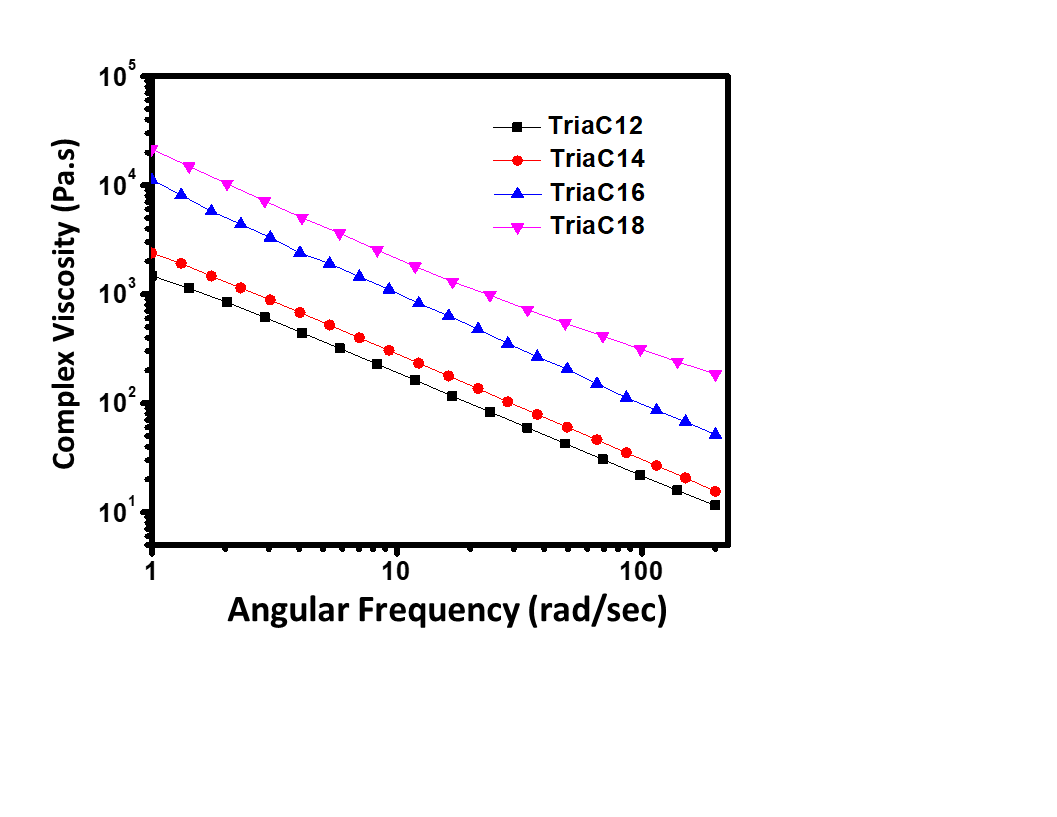

**Figure S17.** Complex viscosity of hydrogelators

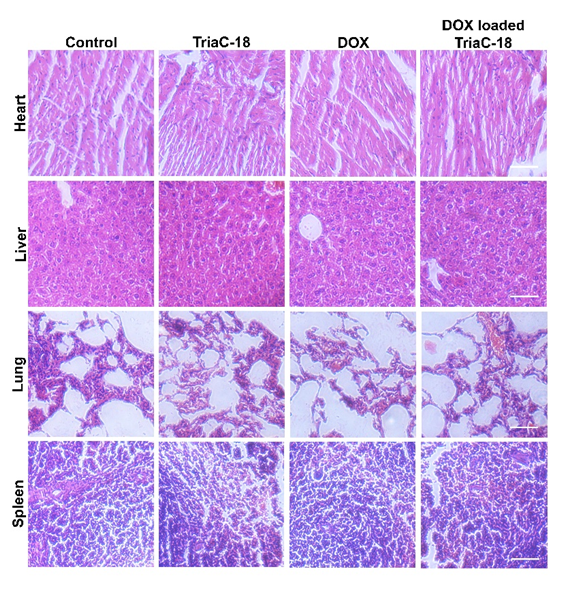

**Figure S18.** H & E staining of different organs (heart, liver, lung and spleen) isolated from different groups of mice

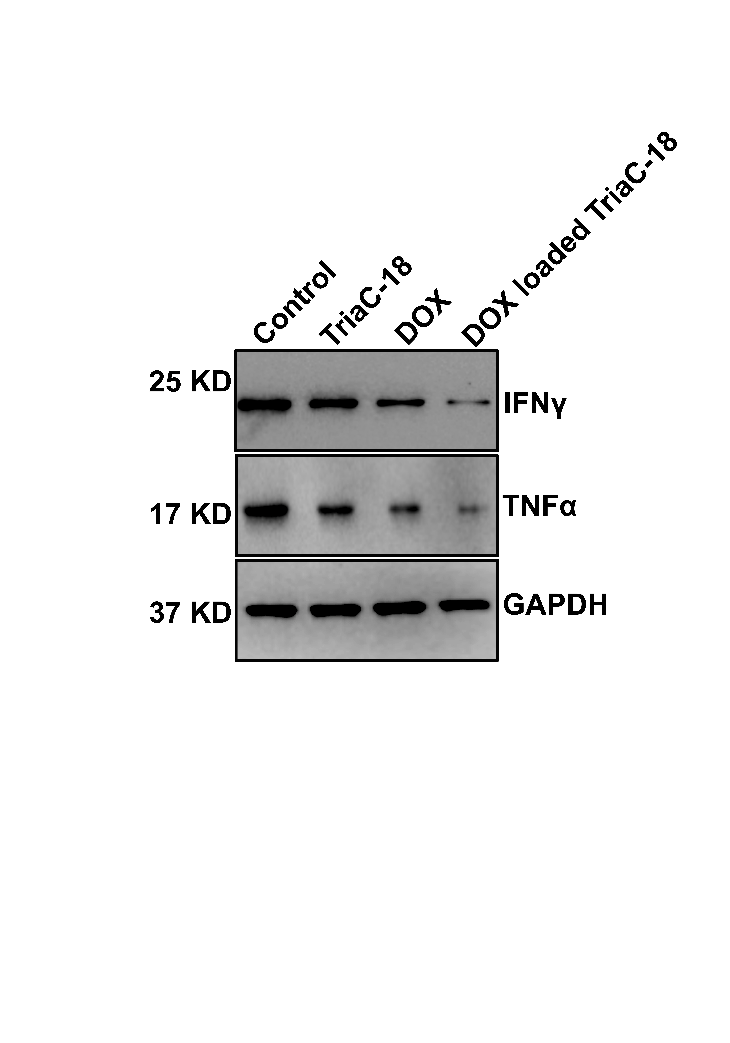

**Figure S19.** Western blot analysis of pro-inflammatory markers of tumor samples
